# *Dyrk1a* gene dosage in glutamatergic neurons has key effects in cognitive deficits observed in mouse models of MRD7 and Down syndrome

**DOI:** 10.1101/2021.05.01.442242

**Authors:** Véronique Brault, Thu Lan Nguyen, Javier Flores-Gutiérrez, Giovanni Iacono, Marie-Christine Birling, Valérie Lalanne, Hamid Méziane, Antigoni Manousopoulou, Guillaume Pavlovic, Loïc Lindner, Mohammed Selloum, Tania Sorg, Eugene Yu, Spiros D. Garbis, Yann Hérault

**Author notes:** Current Address: Proteas Bioanalytics Inc., BioLabs at The Lundquist Institute, Torrance, 90502 CA, USA. **Corresponding author: Yann Hérault**.

## Abstract

Perturbation of the excitation/inhibition (E/I) balance leads to neurodevelopmental diseases including to autism spectrum disorders, intellectual disability, and epilepsy. Mutation in the *DYRK1A* gene located on human chromosome 21 (Hsa21) leads to an intellectual disability syndrome associated with microcephaly, epilepsy, and autistic troubles (MRD7). Overexpression of DYRK1A, on the other hand, has been linked with learning and memory defects observed in people with Down syndrome (DS). *Dyrk1a* is expressed in both glutamatergic and GABAergic neurons, but its impact on each neuronal population has not yet been elucidated. Here we investigated the impact of *Dyrk1a* gene copy number variation in glutamatergic neurons using a conditional knockout allele of *Dyrk1a* crossed with the Tg(Camk2-Cre)4Gsc transgenic mouse. We explored this genetic modification in homozygotes, heterozygotes and combined with the Dp(16*Lipi-Zbtb21*)1Yey trisomic mouse model to unravel the consequence of *Dyrk1a* dosage from 0 to 3, to understand its role in normal physiology, and in MRD7 and DS. Overall, *Dyrk1a* dosage in glutamatergic neurons did not impact locomotor activity, working memory or epileptic susceptibility, but revealed that *Dyrk1a* is involved in long-term explicit memory. Molecular analyses pointed at a deregulation of transcriptional activity through immediate early genes and a role of DYRK1A at the glutamatergic post-synapse by deregulating and interacting with key post-synaptic proteins implicated in mechanism leading to long-term enhanced synaptic plasticity. Altogether, our work gives important information to understand the action of DYRK1A inhibitors and have a better therapeutic approach.

**Author summary:** The Dual Specificity Tyrosine Phosphorylation Regulated Kinase 1A, DYRK1A, drives cognitive alterations with increased dose in Down syndrome (DS) or with reduced dose in mental retardation disease 7 (MRD7). Here we report that specific and complete loss of *Dyrk1a* in glutamatergic neurons induced a range of specific cognitive phenotypes and alter the expression of genes involved in neurotransmission in the hippocampus. We further explored the consequences of *Dyrk1a* dosage in glutamatergic neurons on the cognitive phenotypes observed respectively in MRD7 and DS mouse models and we found specific roles in long-term explicit memory with no impact on motor activity, short-term working memory, and susceptibility to epilepsy. Then we demonstrated that DYRK1A is a component of the glutamatergic post-synapse and interacts with several component such as NR2B and PSD95. Altogether our work describes a new role of DYRK1A at the glutamatergic synapse that must be considered to understand the consequence of treatment targeting DYRK1A in disease.

## Introduction

Down syndrome (DS; Trisomy 21), is the first genetic cause of mental retardation. The 21q22 contains a critical region (named DSCR for “Down syndrome Critical Region”) associated with most DS features including mental retardation. Among genes present in this region, the Dual-specificity Tyrosine-(Y)-phosphorylation-Regulated Kinase 1A (*DYRK1A*), the mammalian homologue of the *Drosophila* minibrain (*mnb*) gene that is essential for normal neurogenesis, is a target for improvement of DS cognition [1]. In addition, Mental Retardation Disease 7 (MRD7) is caused by mutations, including loss-of-function, in *DYRK1A* [2–7], making this gene a critical dosage-sensitive gene for cognitive phenotypes.

The rodent *Dyrk1a* is expressed in foetal and adult brains in dividing neuronal progenitors and later in the adult cerebellum, olfactory bulb and hippocampus. DYRK1A is a serine/threonine kinase with substrates including transcription (CREB, NFAT, STAT3, FKRH), translation (ElF2Be) and splicing factors (SF2, SF3), protein regulating cell cycle (Cyclin B1 and B2) and apoptosis (Caspase 9, P53), synaptic proteins involved in endocytosis/exocytosis, intracellular trafficking, dynamic of the actin cytoskeleton (N-WASP), microtubule formation (MAP1B) and proteins implicated in inter cellular communication (NOTCH, GSK3b). Roles of DYRK1A have been revealed in brain development and neuronal differentiation via the control of critical signalling pathways such as AKT, MAPK/ERK and STAT3 or in synaptic function via the NFAT pathway [8, 9]. Transgenic mice with either excess or haploinsufficiency of *Dyrk1a* show cognitive deficits like those observed in patients with specific impairment of hippocampal-dependent learning and memory [10–12].

Among the mechanisms proposed to underlie the cognitive deficits in DS is glutamatergic and GABAergic neurotransmitter dysfunction. Studies of the DS mouse model Ts65Dn trisomic for about 56% of the human chromosome 21 (Hsa21) syntenic region on mouse chromosome 16 (Mmu16) have revealed excess GABAergic input leading to reduced activation of NMDA receptors and reduction of long-term potentiation (LTP) in the hippocampal CA1 and dentate gyrus (DG) areas [13]. In addition, enhanced hippocampal long-term depression (LTD) has also been observed in the hippocampi of Ts65Dn mice in response to sustained activation of excitatory synapses and attributed to excessive signalling via NMDA receptors [14–16]. Recent evidence supports the contribution of DYRK1A to changes in glutamatergic neurotransmission, with a BAC transgenic mouse line overexpressing *Dyrk1a*, showing alterations in glutamatergic synaptic proteins and normalization of *Dyrk1a* in Ts65Dn mice improving synaptic plasticity, GABAergic/glutamatergic balance, learning and memory [17, 18]. *Dyrk1a* heterozygous knockout mice also present a reduction in the dendritic arborisation and the spine density of glutamatergic pyramidal neurons of the cerebral cortex and alterations in glutamatergic and GABAergic synaptic proteins.

In this context, we hypothesized that change in the dosage of *Dyrk1a* in glutamatergic neurons of the hippocampus and cortex of DS and MRD7 mouse models somehow alter their development and/or normal working in adult brain, leading to the cognitive deficits observed in DS or *Dyrk1a* haploinsufficiency models. Analysis of DYRK1A function in glutamatergic neurons using a knockout approach is not possible as its full KO is homozygote lethal [19]. Thus, we decided to change the gene dosage of *Dyrk1a* in glutamatergic neurons either in a disomic (inactivation of one or two copies of *Dyrk1a*) or trisomic (going back to two copies of *Dyrk1a*) context. We selected the Tg(Camk2a-Cre) transgene to target the Cre recombinase in glutamatergic neurons within the forebrain [20] and we used the Dp(16)1Yey trisomic mouse model (abbreviated as Dp1Yey) containing a segmental duplication of the 22.9 Mb *Lipi-Zfp295* region including *Dyrk1a*, to return to two copies of *Dyrk1a* in the glutamatergic neurons of the Dp1Yey. This model has the advantages to include 65% of Hsa21 mouse gene orthologs and to be devoid of the 50 DS-irrelevant trisomic genes that are present on the Ts65Dn mini chromosome [21]. Dp1Yey mice present defects in working memory, long-term episodic memory, and associative learning. In addition to those tests, we also tested the impact of *Dyrk1a* gene dosage on the mouse social behaviour as MRD7 patients display autistic traits.

## Results

### *Dyrk1a* is expressed in Camk2a-positive cells and its full inactivation in the glutamatergic neurons induces brain defects

*Dyrk1a* is ubiquitously expressed in different neuronal cell populations of the brain but with regional differences: the protein level being higher in the olfactory bulb, cerebellar cortex, cortical structures and granular and pyramidal cell layers of the hippocampus [22]. We checked DYRK1A expression in adult glutamatergic neurons, by co-immunohistochemical localisation with an antibody against CAMK2A and DYRK1A. In the wild-type adult mouse, both proteins were found in pyramidal and granular neurons of the hippocampus and dentate gyrus and in neurons of the cortex (Figure 1A).

**Figure 1:**
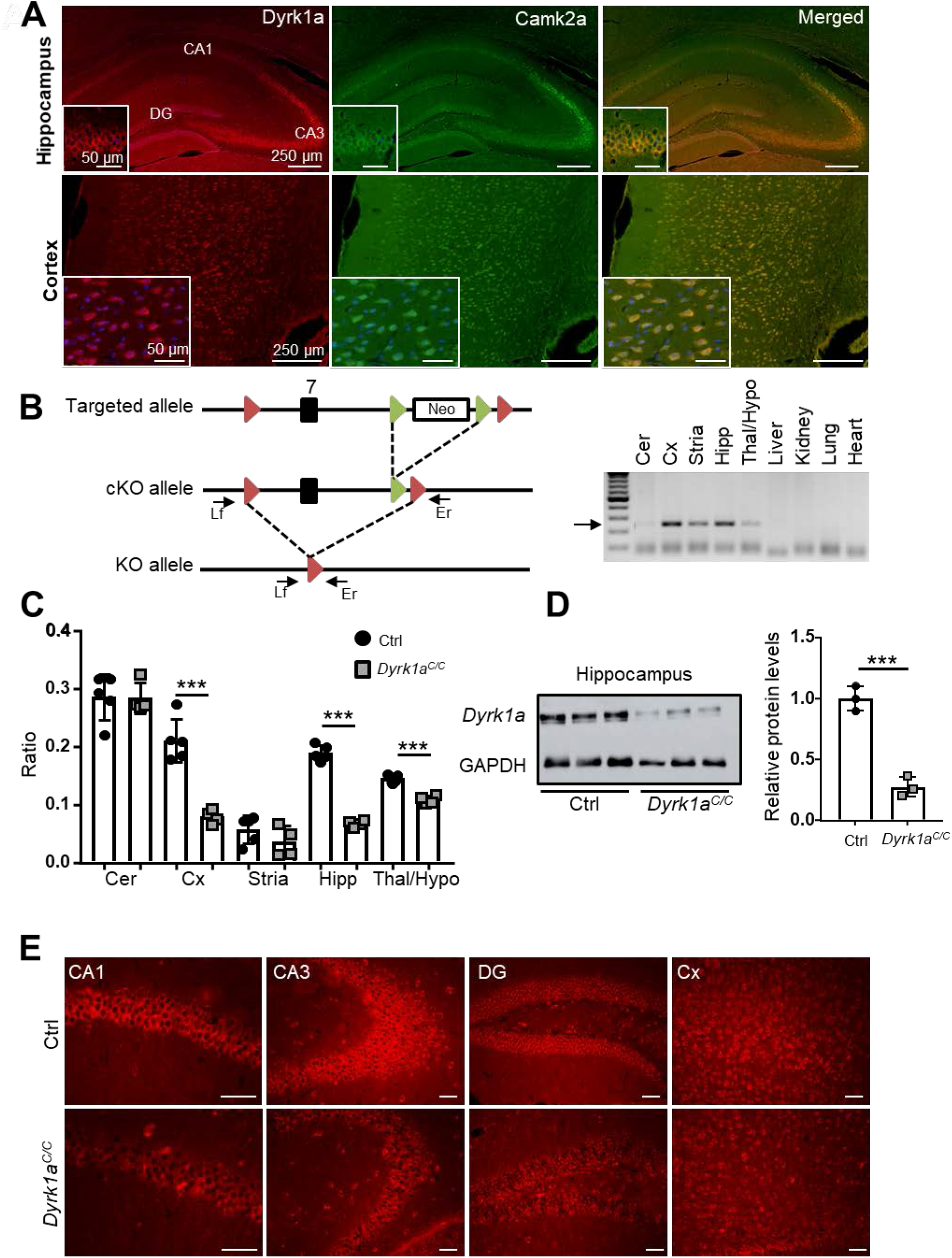
Generation of mice deficient for Dyrk1a in the glutamatergic neurons. (A) DYRK1A (red) co-localizes with CAMK2A (green) in the glutamatergic pyramidal neurons of the CA1-3, the granular neurons of the dentate gyrus (DG) and in the cortex. (B) Targeting strategy for conditional inactivation of *Dyrk1a*. Exon 7 containing the serine/threonine protein kinase active site was flanked with loxP sites (red arrowheads) in two steps: a targeted allele was first generated by homologous recombination in ES cells, then *in vivo* expression of the Flp recombinase resulted in recombinaison of the FRT sites (green arrowheads) and removal of the selection cassette (white box) generating the conditional allele (cKO). The knock-out allele (KO) was observed in the brain of *Dyrk1a^C/C^*. Arrows represent primers for PCR genotyping. Genomic DNA was isolated from different organs from a *Dyrk1a^C/C^* mouse and genotyped for the presence of the knock-out allele with primers Lf and Er, giving a 232 bp PCR product for the KO allele. (C) Ratio of relative mRNA of *Dyrk1a* in different brain structures in *Dyrk1a^C/C^* and disomic control mice. (D) Autoradiographic image and quantification of immunoblots of DYRK1A protein in the hippocampus of *Dyrk1a^C/C^* mice relative to control mice. Band intensities were estimated using ImageJ and normalized against the loading control GAPDH. (E) DYRK1A immunohistochemistry of coronal brain sections at the level of the hippocampus from control and *Dyrk1a^C/C^* mice. Data are presented as point plots with mean ± SD with unpaired Student’s t-test, *p<0.05, **p<0.01, ***p<0.001 (n=5 ctrl and 4 *Dyrk1a^C/C^* hippocampus for mRNA analysis and n=3 per genotype for protein analysis). CA1, Cornus Ammonis 1; CA3, Cornus Ammonis 3; DG, Dentate Gyrus; Cer: Cerebellum; Cx, Cortex.

To better understand the function of DYRK1A in glutamatergic neurons we inactivated both copies of *Dyrk1a* using a conditional approach, to generate a full knock-out in those neurons. A floxed *Dyrk1a* allele (*Dyrk1a^cKO^* allele) was designed such that exon 7 that codes for the serine/threonine protein kinase active site signature domain was flanked by two *loxP* sites (Figure 1B). We used the Tg(Camk2aCre)4Gsc transgene [20] to generate the *Dyrk1a^Camk2aCre^* allele (shortened as *Dyrk1a^C^*) and we checked the ability of the Cre to recombine the *Dyrk1a* floxed allele in *Dyrk1a^Camk2aCre/Camk2aCre^* (recombination of both *Dyrk1a^cKO^* alleles with the Cre recombinase; noted here *Dyrk1a^C/C^*) mice. The generation of the deleted allele was detected by PCR analysis exclusively in brain areas where *Camk2a* is expressed (Figure 1B). Quantification of *Dyrk1a* mRNA in different brain regions confirmed that *Dyrk1a* is expressed at different relative levels in brain subregions (Figure 1C). Nevertheless, decrease of the *Dyrk1a* transcripts was found in the hippocampus, cortex and thalamus/hypophysis but not in the cerebellum of *Dyrk1a^C/C^* mice (Figure 1C). Loss of the DYRK1A protein was confirmed in the hippocampus by Western blot analysis (Figure 1D) and immunohistology (Figure 1E). This reduction was more evident within the pyramidal cell layers of the CA1 and CA3 composed mostly of glutamatergic neurons.

We analysed the implication of DYRK1A in glutamatergic neurons by looking at brain morphology and cognitive phenotypes. Brain weight was significantly decreased in *Dyrk1a^C/C^* mice compared to control mice (90% of the control weight; Figure 2A). Morphometric analysis at Bregma −1.5 (Figure 2B) unravelled reduced surface area of the total brain surface in *Dyrk1a^C/C^* mice (~88% of control; Figure 2C). The area of the hippocampus including the cornus ammonis fields (CA1, CA2 and CA3) and dentate gyrus (DG) did not significantly differ between the two genotypes (Figure 2D). We measured the thickness of the oriens layer at the CA1, CA2 and CA3 levels, of the pyramidal layer at the CA1 level, of the radiatum layer, of the CA1 and DG molecular layers and of the granular layer of the DG and did not find any difference between control and *Dyrk1a^C/C^* mice (Supplementary figure 1A). We also counted neurons within the CA1 and did not find any modification in the density of pyramidal neurons (Supplementary figure 1B). Specific decrease in cortical thickness was observed at the level of the dorsal motor cortex (~78% of controls, Figure 2E) and of the somatosensory cortex (~76% of controls, Figure 2E) whereas decrease in thickness at the more ventral auditory cortex level was not significant (Fig. 2E). Measurements of the thickness of different layers in the somatosensory cortex (Figure 2F) indicate decrease in the thickness of molecular layer I, external granular and pyramidal layers II/III, internal pyramidal layer V and internal polymorphic layer VI (Figure 2G). Only the internal granular layer IV was found unchanged (Figure 2G). To investigate how change in cellularity might relate to cortical thickness, we counted the number of cells present in SSC layers II-III, V and VI. We found that cellular density in layers II-III, V and VI was increased by about 30% in *Dyrk1a^C/C^* mice (Figure 2H) with the total number of neurons unchanged between *Dyrk1a^C/C^* and controls, suggesting an impact of *Dyrk1a* inactivation on cell morphology or tissue organization.

**Figure 2:**
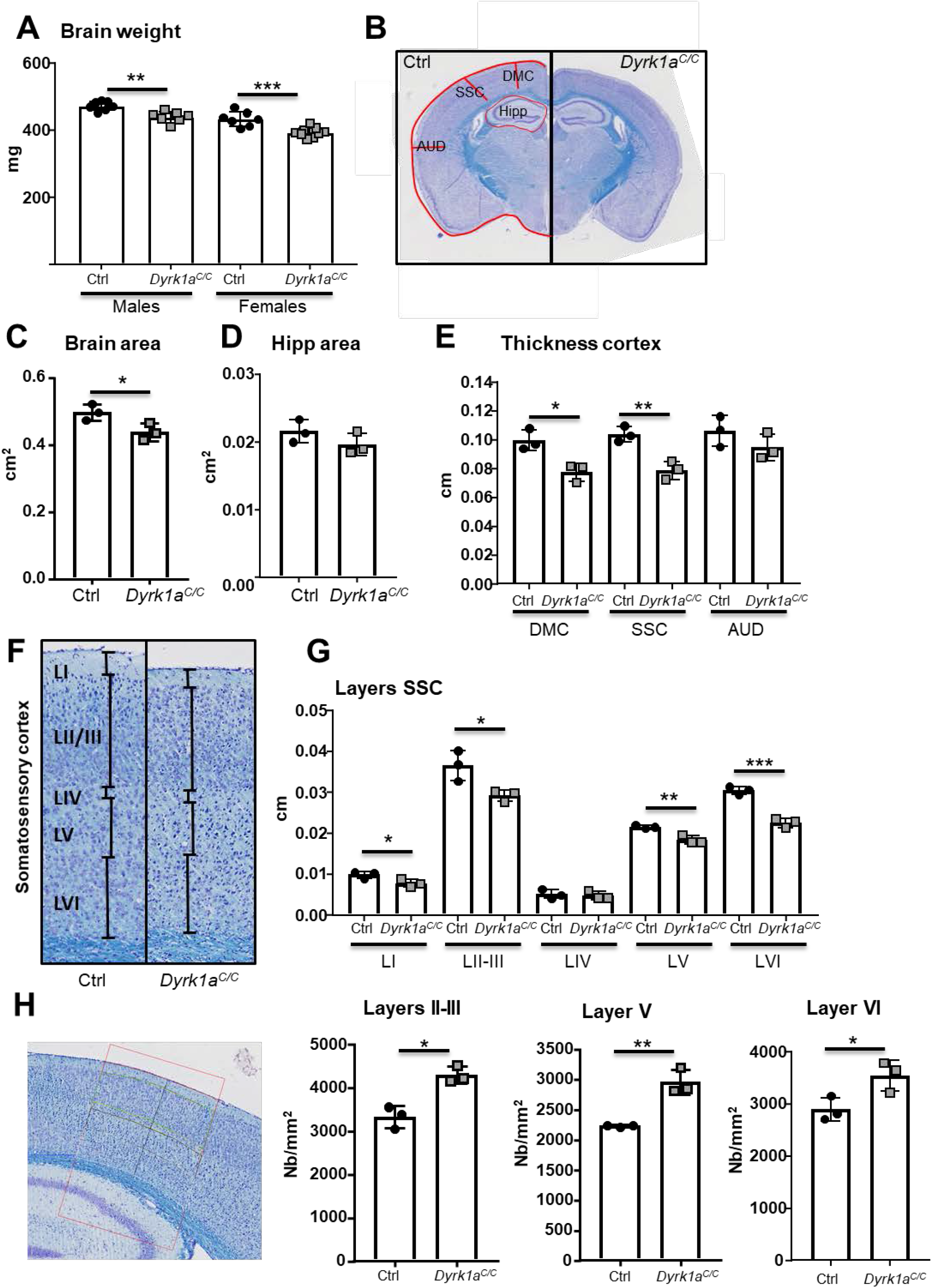
Consequence of *Dyrk1a* inactivation in glutamatergic neurons on brain morphology. (A) Brain weight from male and female mice aged 3 months old (n=7-9 per genotype). (B) Representative coronal sections of control (Ctrl) (left) and *Dyrk1a^C/C^* (right) brains at Bregma −1.5 stained with cresyl violet and luxol blue that were used for measurements (Magnification 20X). (C) Dot plots of total brain area measurements (red line around the brain in B). (D) Dot plots of hippocampal areas (red area around hipp in B). (E) Measurements of the thickness of the cortex at the 3 levels represented by red lines in figure B. (F) Representative cresyl violet and luxol blue stained coronal sections of somatosensory cortex layers in control (ctrl) and *Dyrk1a^C/C^* brains at Bregma −1.5. (G) Measurements of the thickness of the different layers presented in figure F. (H) Relative density of cells counted in layers II-III, V and VI within a frame of 0.1 cm width (see figure) at the level of the somatosensory cortex. Data are presented as point plots with mean ± SD with unpaired Student’s t-test, *p<0.05, **p<0.01, ***p<0.001 (n=3 females per genotype). AUD: auditory cortex, SSC: somatosensory cortex, DMC: dorso motor cortex, Hipp: hippocampus.

### Full *Dyrk1a* inactivation in the glutamatergic neurons impacts general behaviour and cognition

To analyse mouse behaviour in *Dyrk1a^C/C^* mice, we first focused our attention on locomotor activity and exploratory activity. Measurement of horizontal, or vertical, locomotor activity during circadian cycle did not differ in *Dyrk1a^C/C^* mice compared to control mice (Supplementary figure 2A, B). The analysis of exploratory behaviour in a novel environment (open field (OF) test) indicated normal locomotor activity for the *Dyrk1a^C/C^* mice (Supplementary figure 2C) but their exploratory pattern was altered as they spent significantly more time in the centre of the OF (Figure 3A), suggesting a decreased anxiety. This phenotype was confirmed in the elevated plus maze with *Dyrk1a^C/C^* mice spending significantly more time in the open arms than control mice (Figure 3B). In this test, *Dyrk1a^C/C^* mice were also more active, visiting more arms than the control mice (Figure 3C). The locomotor performance was assessed in the rotarod task. *Dyrk1a^C/C^* mice exhibited slightly better performance in this test than their control littermates, with an increase in latency to fall, indicating that motor balance is not affected in those mice (Figure 3D).

**Figure 3:**
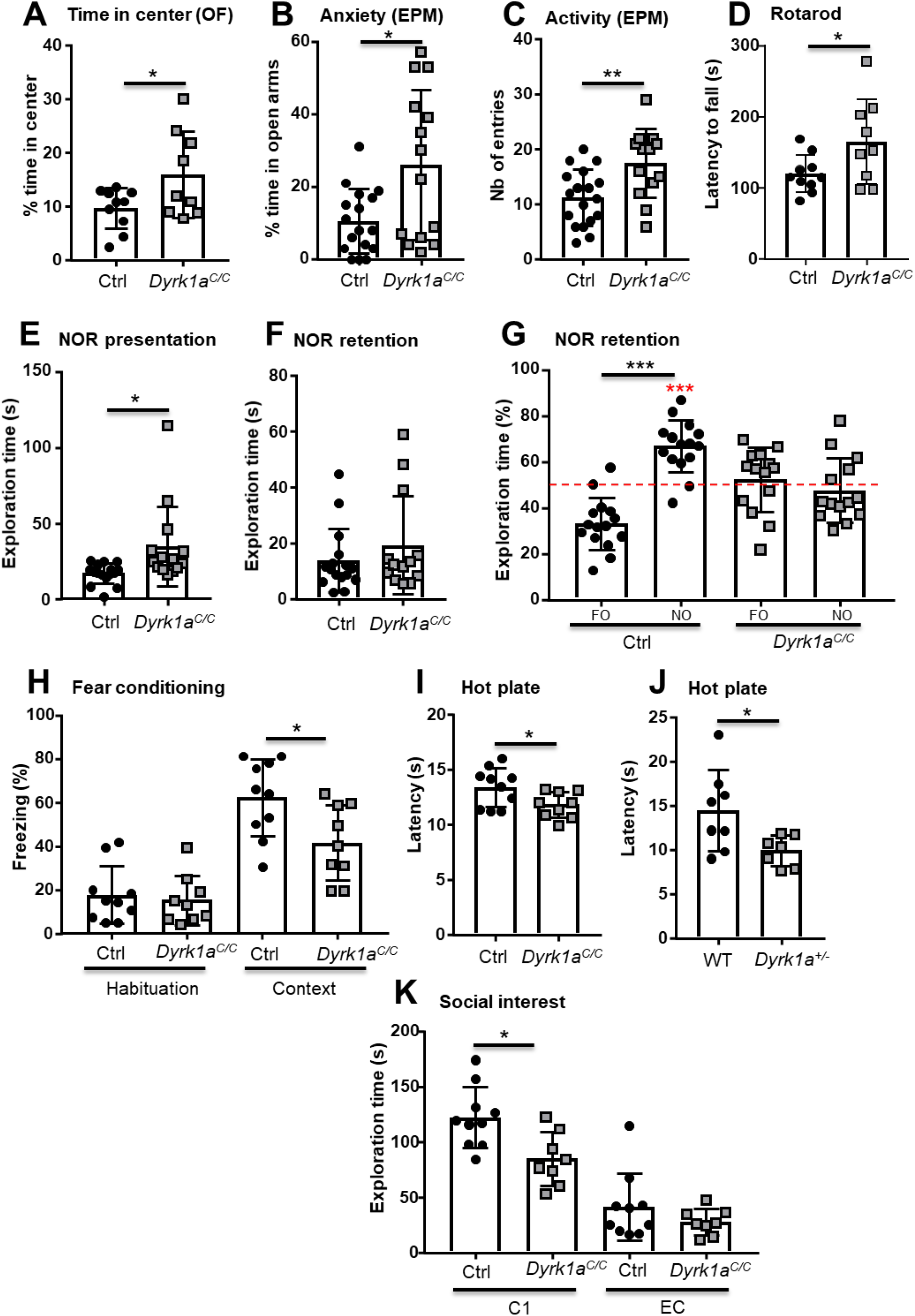
Impact of *Dyrk1a* inactivation in glutamatergic neurons on general behavior, locomotor activity and cognition. (A) The exploratory behavior of a new environment was analyzed by the percentage of time spent in the center of an open field over 30 min of test. *Dyrk1a^C/C^* mice spent more time in the center of the arena, suggesting that they are less anxious. (B) Confirmation of the phenotype in a new group of mice using the Elevated Plus Maze test (EPM) with *Dyrk1a^C/C^* mice showing a higher percentage of time spent in the open arms of the maze. (C) Mouse activity, measured by the number of entries in both open and closed arms, was increased in *Dyrk1a^C/C^* animals. (D) Evaluation of the locomotor performance on the rotarod during consecutive trials with increased rotational speeds. The latency is the mean of 3 independent trials. *Dyrk1a^C/C^* mice showed increased performance. (E) In the NOR test, the percentage of time spent exploring the familiar (FO) and the new (NO) objects show that control mice spend significantly more time on the NO while *Dyrk1a^C/C^* mice do not make any difference between the two objects. (F) Exploration times of the two identical objects during the presentation phase of the NOR show that *Dyrk1a^C/C^* mice tend to have an increased exploration time. (G) Total exploratory time of the two objects during the test indicates that the absence of object discrimination of the *Dyrk1a^C/C^* mice is not due to a lack of interest of the objects. (H) Percentage of freezing time during the habituation phase (before the foot shock; basal level of activity) (Habituation; Mann-Whitney rank sum test, p=0.71) and during the 6 min of contextual exposure 24 hours later indicate a deficit of contextual learning in *Dyrk1a^C/C^* mice. (I-J) Pain sensitivity was evaluated by measuring the mouse latency to elicit a response to pain when put on a plate (52°C). In this test, *Dyrk1a^C/C^* and *Dyrk1a^+/−^* mice had a lower threshold than their control littermates. (K) Interest in social interaction was measured by the time spent sniffing the cage containing a congener (C1) during the Crawley test. This time was reduced for *Dyrk1a^C/C^* mice compared to control mice (unpaired t-test p=0.01) while the time spent exploring the empty cage (EC) was the same between the two groups (unpaired t-test p=0.23). Data are presented as point plots with mean ± SD. Statistical analyses were done with unpaired Student’s t-test or Mann-Whitney rank sum test if normality test failed, except E: paired T-test FO vs NO and one sample T-test vs 50% mean (in red: ***p<0.001); n=8-10 per genotype. B-C (EPM) and E-G (NOR) were done with another batch of mice, n=14-15 per genotype; *p<0.05, **p<0.01, ***p<0.001.

Impact of the loss of *Dyrk1a* in glutamatergic neurons on cognition was evaluated using different memory tests. Working memory was assessed by recording spontaneous alternation in the Y-maze. The percentage of alternation between the three arms was similar between *Dyrk1a^C/C^* and control mice (Supplementary figure 2D) indicating a normal working memory in both genotypes. In this test, the number of visited arms during the 5 min session was not significantly different in the *Dyrk1a^C/C^* mice compared to the controls, although those mice showed more variability (Supplementary figure 2E). Long-term explicit memory requiring the hippocampus and related medial and temporal lobe structures was tested with the novel object recognition test (NOR) with 24 hours delay. Although *Dyrk1a^C/C^* mice showed as much interest exploring the objects during the presentation session (Figure 3E) and during the discrimination session (Figure 3F) compared to control animals, they did not make any difference between the two objects during the retention trial (Figure 3G) by contrast to control mice who spent significantly more time on the novel object compared to the familiar one. Thus, the NOR test unravelled a deficit in long-term explicit memory in the *Dyrk1a^C/C^* mice. Then, we tested associative learning using the contextual fear-conditioning test. During the habituation, mice showed the same basal level of freezing whatever their genotypes were (Figure 3H, Habituation). However, *Dyrk1a^C/C^* mice showed significantly less freezing than control mice during contextual discrimination, indicating poorer performance in contextual learning (Figure 3H, Context). During the cued learning, *Dyrk1a^C/C^* mice responded like control mice to the conditioned stimulus, indicating normal cued fear (Supplementary figure 2F). As decreased freezing could be due to a deficit in pain sensitivity rather than a deficit in memory, we tested the mice in the hot plate test. *Dyrk1a^C/C^* mice had a decreased latency to elicit a first response to noxious thermal stimulus, suggesting that they were more sensitive to pain than control mice (Figure 3I). As pain sensitivity was never tested in *Dyrk1a* knock-out heterozygous mice (shortened as *Dyrk1a^+/−^*), we also tested those mice in the hot plate. We also found that those mice are more sensitive to pain (Figure 3J).

As in Human *DYRK1A* heterozygous mutations lead to autistic behaviour in MRD7, mouse sociability was investigated in this full inactivation of *Dyrk1a* in the glutamatergic neurons. We presented an empty cage and a cage containing a congener to the tested mouse and measured the time spent by the tested mouse to sniff either cage. Both *Dyrk1a^C/C^* and control mice showed social preference as they spent significantly more time sniffing the cage containing the congener than the empty cage (Supplementary figure 2G, Social preference). However, the total amount of time spent with their congener was decreased in *Dyrk1a^C/C^* mice compared to control mice whereas the time spent exploring the empty cage did not differ (Figure 3K). Preference for social novelty was tested by placing a new congener in the empty cage. Both genotypes spent significantly more time sniffing the cage containing the new congener compared to the cage with the familiar one (>60% of the time allocated for the new congener) (Supplementary figure 2H, Social novelty preference). There was also no significant difference in the total time control and transgenic animals spent sniffing both congeners (Supplementary figure 2I, Social contact).

Finally, as *DYRK1A* haploinsufficiency in human is causing epilepsy, we challenged the homozygous inactivation in *Dyrk1a^C/C^* and control mice with two different doses of the seizure-provoking agent pentylenetetrazol (PTZ) and the occurrence of myoclonic, clonic and tonic seizures was scored. At both 30 mg/kg (Supplementary figure 2J and 50 mg/kg (Supplementary figure 2K), *Dyrk1a^C/C^* susceptibility to seizure was similar to control mice. Altogether, those results indicate that *Dyrk1a* full inactivation in glutamatergic neurons does not increase susceptibility to PTZ-induced seizure.

Hence, *Dyrk1a* inactivation in glutamatergic neurons only impacts specific cognitive function such as explicit long-term memory, contextual fear memory and exploratory behavior while having no impact in others such as working memory, social behaviour and epileptic susceptibility.

### *Dyrk1a* inactivation in the glutamatergic neurons lowers expression of genes involved in neurotransmission in the hippocampus, while enhancing expression of genes implicated in the regulation of transcription

The hippocampus is a key structure in memory formation. Long-term object recognition memory analysed in the NOR test was shown to require interaction between the hippocampus and the perirhinal cortex [23–25] while contextual fear memory involves a neural circuit including the hippocampus, amygdala and medial prefrontal cortex [26]. Although we could not detect any morphological defect in the hippocampus of *Dyrk1a^C/C^* mice, those mice are defective in both long-term recognition and contextual fear memories. To unravel the potential molecular mechanisms underlying the learning defects of *Dyrk1a^C/C^* mice, we performed genome-wide transcriptional profiling (RNA-seq) of *Dyrk1a^C/C^* and control mice in the hippocampus at postnatal day 30. Analysis of the RNA-seq exon reads (DEseq algorithm, P<0.025) identified 297 up-regulated and 257 down-regulated genes in *Dyrk1a^C/C^* compared with controls (Supplementary Table S1,). To determine the putative cell types associated with the deregulated genes, we compared the sets of up- and down-regulated genes with the markers of hippocampal cell types obtained from single cell RNA-seq [PMID: 25700174][PMID: 29273784] (see methods). As a result, up-regulated genes were predominantly enriched in oligodendrocyte-expressed genes (hypergeometric test, *bonferroni* corrected, P<3.4E-13) whereas down-regulated genes were enriched in neuronal markers (hypergeometric test, *bonferroni* corrected, pyramidal markers P<2.3E-4, interneuronal markers P<8.3E-3) (Supplementary Table S2). This decrease in neuronal markers expression is not reflected by a decrease in neuronal cells in the hippocampus as we did not observe a decrease in the thickness of the pyramidal cell layers or the DG granular cell layer, and did not find a deficit in the number of neurons within the CA1 of the *Dyrk1a^C/C^* hippocampus (Supplementary figure 1). Next, we performed GO enrichment analyses of the lists of up- and down-regulated genes with using a *Benjamini* cut-off of P < 0.05 (Table 1). The strongest enrichments for up-regulated genes were related to transcriptional regulation and DNA methylation. This category of genes did not have any overlap with the oligodendrocyte overexpressed genes at the exception of the SRY-related HMG-box transcription factor *Sox8* and we did not find any enriched specific function for the list of the oligodendrocyte markers that are up-regulated in *Dyrk1a^C/C^* hippocampi. We counted the number of Olig2+ cells in the corpus-callosum and found no difference between *Dyrk1a^C/C^* and control mice, suggesting that increased oligodendrocyte markers is not due to an increased number of oligodendrocytes (Supplementary figure 3). Interestingly, among up-regulated genes, we found *Nr4a1 (Nurr77), Arc (Arg3.1), Npas4, Fos (cFos), Egr1 (Zif268) and Fosb*, six immediate-early genes (IEGs) encoding proteins involved in transduction signals that are induced in response to a wide variety of cellular stimuli and that are implicated in neuronal plasticity. Looking at known late response genes known to be activated by NPAS4 [27] in glutamatergic neurons, only three out of thirty-four (10%) of them were significantly deregulated in the hippocampus of *Dyrk1a^C/C^* compared to control mice (Supplementary Table S3), with *Fam198b* being up-regulated and *Csrnp1* and *Slc2a1* being down-regulated. Among target genes of NPAS4 shared between excitatory and inhibitory cells, four out of twelve (~30%) that we looked at were found deregulated in the hippocampus of *Dyrk1a^C/C^* compared to control mice (*Lmo2* and *Fosl2*, up-regulated; *Mylk* and *Nptx2*, down-regulated). Down-regulated genes found in the hippocampus transcriptome of *Dyrk1a^C/C^* mice were associated with presynaptic vesicle exocytosis, regulation of neurotransmitter levels and neuron projection, and pointed at a perturbation of chemical synaptic transmission via the deregulation of proteins involved in synaptic vesicle exocytosis. Particularly, genes coding for proteins of the SNARE complex (*Snap25, Stx1a, Napa* and *Napb*), regulating its activity (*Doc2b, Snph*) or implicated in vesicular synaptic cycle (*Anxa7, Amph, Syn2, Syngr1*) were found down-regulated in the hippocampus of *Dyrk1a^C/C^* mice. This complex is known to mediate synaptic vesicle docking and fusion with the presynaptic membrane during neuromediator release. The SNARE complex was recently found, also with NPAS4, as a common pathway misregulated in models of DS overexpressing DYRK1A (Duchon et al, HMG).

**Table 1:**
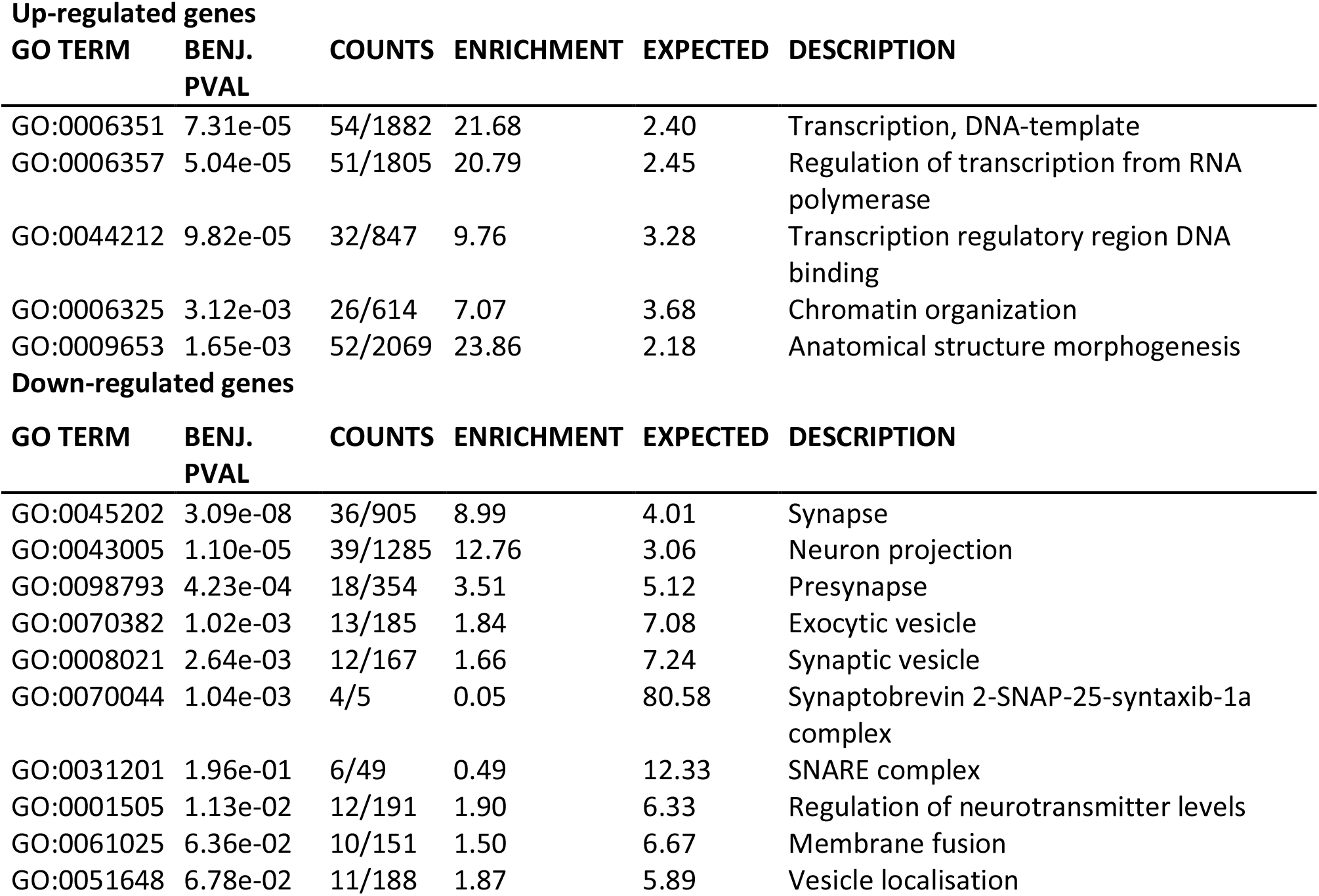
GO enrichment analyses of the up- and down-regulated genes expressed in the *Dyrk1a^C/C^* hippocampus (*Benjamini* cut-off of P < 0.05).

| Up-regulated genes |  |  |  |  |  |
| --- | --- | --- | --- | --- | --- |
| GO TERM | BENJ. PVAL | COUNTS | ENRICHMENT | EXPECTED | DESCRIPTION |
| GO:0006351 | 7.31e-05 | 54/1882 | 21.68 | 2.40 | Transcription, DNA-template |
| GO:0006357 | 5.04e-05 | 51/1805 | 20.79 | 2.45 | Regulation of transcription from RNA polymerase |
| GO:0044212 | 9.82e-05 | 32/847 | 9.76 | 3.28 | Transcription regulatory region DNA binding |
| GO:0006325 | 3.12e-03 | 26/614 | 7.07 | 3.68 | Chromatin organization |
| GO:0009653 | 1.65e-03 | 52/2069 | 23.86 | 2.18 | Anatomical structure morphogenesis |
| Down-regulated genes |  |  |  |  |  |
| GO TERM | BENJ. PVAL | COUNTS | ENRICHMENT | EXPECTED | DESCRIPTION |
| GO:0045202 | 3.09e-08 | 36/905 | 8.99 | 4.01 | Synapse |
| GO:0043005 | 1.10e-05 | 39/1285 | 12.76 | 3.06 | Neuron projection |
| GO:0098793 | 4.23e-04 | 18/354 | 3.51 | 5.12 | Presynapse |
| GO:0070382 | 1.02e-03 | 13/185 | 1.84 | 7.08 | Exocytic vesicle |
| GO:0008021 | 2.64e-03 | 12/167 | 1.66 | 7.24 | Synaptic vesicle |
| GO:0070044 | 1.04e-03 | 4/5 | 0.05 | 80.58 | Synaptobrevin 2-SNAP-25-syntaxin-1a complex |
| GO:0031201 | 1.96e-01 | 6/49 | 0.49 | 12.33 | SNARE complex |
| GO:0001505 | 1.13e-02 | 12/191 | 1.90 | 6.33 | Regulation of neurotransmitter levels |
| GO:0061025 | 6.36e-02 | 10/151 | 1.50 | 6.67 | Membrane fusion |
| GO:0051648 | 6.78e-02 | 11/188 | 1.87 | 5.89 | Vesicle localisation |

### Behavioural defects are induced in *Dyrk1a^C/+^* mice while partial rescue of memory alterations is observed in Dp1Yey/*Dyrk1a^C/+^* mice

We investigated the respective consequences of *Dyrk1a* dosage in glutamatergic neurons on the cognitive phenotypes observed respectively in MRD7 and DS mouse models. We analysed mice heterozygous for the *Dyrk1a* knockout allele in glutamatergic neurons to investigate the implication of this gene in the cognitive phenotypes of MRD7. We also performed a rescue experiment consisting on the return to two copies of *Dyrk1a* in the glutamatergic neurons of Dp1Yey trisomic mice. For this, we compared animals carrying *Dyrk1a^Camk2aCre/+^* (noted *Dyrk1a^C/+^*), Dp1Yey, and Dp1Yey/*Dyrk1a^Camk2aCre/+^* (noted Dp1Yey/*Dyrk1a^C/+^*), with *Dyrk1a^cKO/+^* as controls in behavioural tests. Mouse locomotor behaviour was tested in the open-field (OF). *Dyrk1a^C/+^* mice did not show any significant difference in locomotor activity compared to controls. Surprisingly, whereas in our conditions Dp1Yey mice travelled the same total distance in the OF as control mice, Dp1Yey/*Dyrk1a^C/+^* mice travelled significantly more distance (Figure 4A), suggesting that those mice are hyperactive. The percentage of time spent by the mice in the centre of the open field arena did not differ between genotypes (Figure 4B), suggesting normal anxiety-related behaviour. Mice heterozygous for *Dyrk1a* in glutamatergic neurons presented the same behaviour as control mice, indicating that removing only one copy of *Dyrk1a* is not enough to trigger decreased anxiety, as observed in the complete knockout of *Dyrk1a* in glutamatergic neurons (Figure 4B vs Figure 3A). Analysis of working memory was done using the Y maze test. All the four groups of mice visited the same number of arms during the test, suggesting a normal locomotor activity (Supplementary figure 4A). On the other hand, Dp1Yey mice showed lower percentage of spontaneous alternation as compared to control mice (Figure 4C), confirming the phenotype already observed in previous studies [28] (Duchon and Herault, HMG). This decreased performance was not restored by *Dyrk1a* normalization in glutamatergic neurons (Figure 4C). Haploinsufficiency of *Dyrk1a* in those neurons, like the inactivation of the two copies of *Dyrk1a*, did not trigger any change in working memory (Figure 4C). We therefore also tested *Dyrk1a^+/−^* mice in the same test. *Dyrk1a^+/−^* animals showed the same activity (number of visited arms, Supplementary figure 4B) and the same level of alternation as their wild-type littermates, indicating a normal working memory (Supplementary figure 4C).

**Figure 4:**
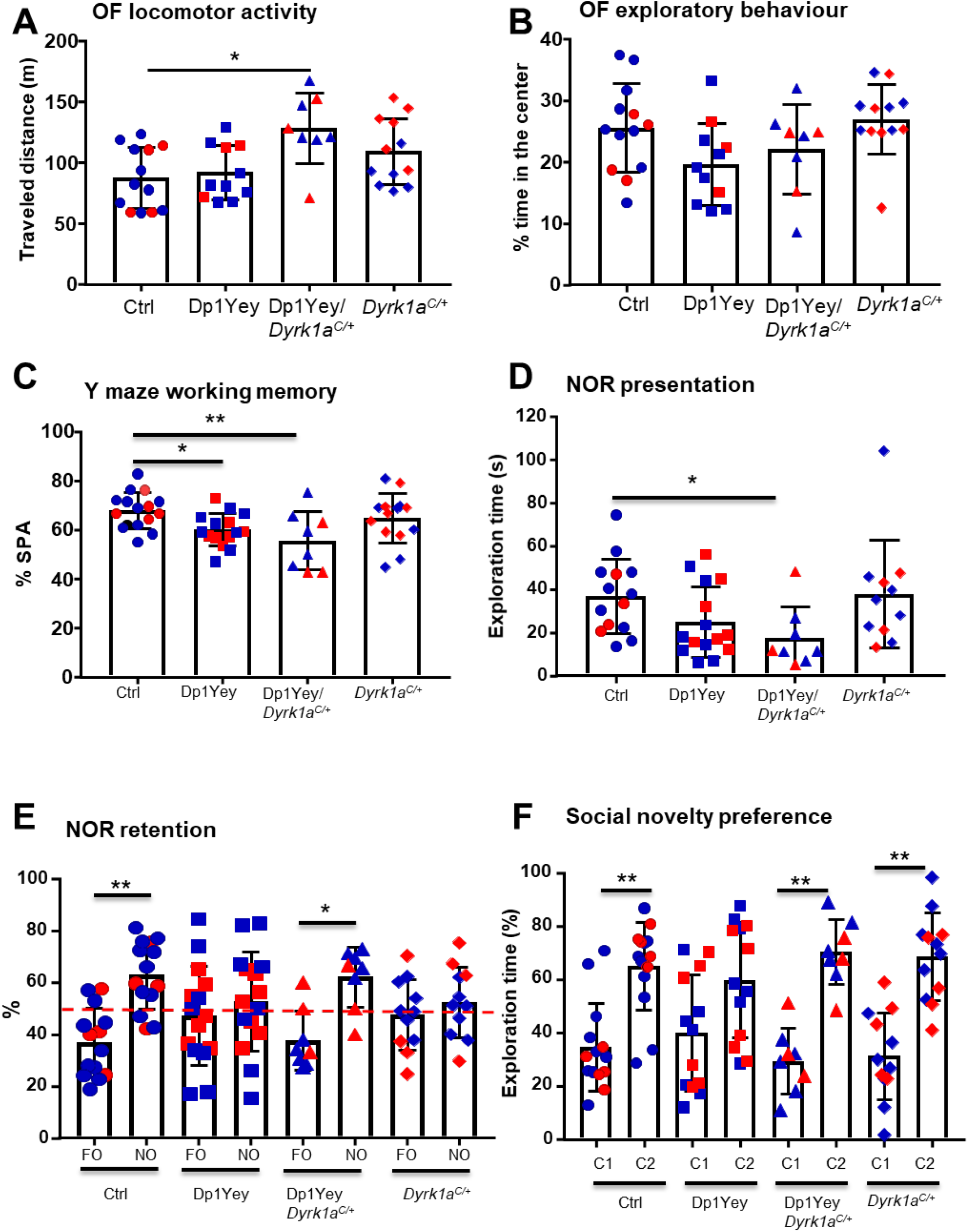
Consequence of the normalization of *Dyrk1a* in glutamatergic neurons of Dp1Yey mice on animal cognition. (A) The total distance travelled in the open field during a 30 min session is significantly increased in the Dp1Yey/*Dyrk1a^C/+^* mice compared to the control mice (Kruskal-Wallis One Way Analysis of Variance on Ranks, *p*=0.009 with Dunn’s post hoc multiple comparison procedures versus control, Dp1Yey/*Dyrk1^C//+^* vs control, *p*=0.007). (B) Percentage time spent in the center of the OF does not vary between genotypes (One way ANOVA, F(3, 40)=2.76, p=0.054). (C) Percentage of spontaneous alternation of the mice during a 5 min session in a Y-maze. Lower percentage of alternation was found in Dp1Yey mice and in Dp1Yey/*Dyrk1a^C/+^* indicating a deficit in working memory in Dp1Yey mice that is not rescued in Dp1Yey/*Dyrk1a^C/+^* mice (One way ANOVA, Holm-Sidak method for multiple comparisons versus control group, F(3,49)=4.3, *p*=0.009, Dp1Yey vs control *q*=2.48, *p*=0.03, Dp1Yey/*Dyrk1a^C/+^ q*=3.24, *p*=0.006). (D-E) Novel object recognition was assessed with 24 hour time laps. (D) Time spent exploring the two identical objects during the first object presentation session was decreased in Dp1Yey/*Dyrk1a^C/+^* mice (Kruskal-Wallis One Way Analysis of Variance on Ranks, *p*=0.02 with Dunn’s post hoc multiple comparison procedures versus control, Dp1Yey/*Dyrk1a^C//+^* vs control, *p*=0.02). (E) When introducing the novel object during the retention period, both control and Dp1Yey/*Dyrk1a^C/+^* lines spent significantly more time exploring the novel object than the familiar one (Paired t-test novel object vs familiar object, ctrl *p*=0.003, Dp1Yey/*Dyrk1a^C/+^ p*=0.02), whereas *Dp1Yey* and *Dyrk1a^C/+^* mice did not (Paired t-test novel object vs familiar object, Dp1Yey *p*=0.57, *Dyrk1a^C/+^ p*=0.56), revealing a significant deficit in memory for both Dp1Yey and *Dyrk1a^C/+^* mice which is rescued in Dp1Yey/*Dyrk1a^C/+^* mice. (F) Mice were tested for social novelty preference. All the genotypes but Dp1Yey spent significantly more time sniffing the new congener (Paired t-test congener vs empty cage, ctrl *p*=0.004, Dp1Yey *p*=0.13, Dp1Yey/*Dyrk1a^C/+^ p*=0.002, *Dyrk1a^C/+^ p*=0.002). Data are presented as point plots with mean ± SD. (n=8-15 per genotype, *p<0.05, **p<0.01) Males (in blue) and females (in red) are pooled in the same graph as the statistical analyses did not reveal significant effect of sex.

We further tested the mice in the NOR test for long term reference memory. Both control and Dp1Yey/*Dyrk1a^C/+^* mice showed a significant preferential exploration of the novel object during the retention trial (Figure 4E) whereas Dp1Yey and *Dyrk1a^C/+^* mice spent the same time on the two objects (Figure 4E). The deficit of novel object exploration during the retention phase in Dp1Yey and *Dyrk1a^C/+^* mice was not due to a lack of familiar object exploration during the presentation phase as both genotypes showed similar exploration times than control mice (Figure 4D). Only Dp1Yey/*Dyrk1a^C/+^* showed a slight decrease in object exploration compared to control mice during the presentation phase mice (Figure 4D), but this did not impair their retention capacity during the test phase. Hence, the deficit in object recognition in Dp1Yey mice could be rescued by normalization of *Dyrk1a* copy number in glutamatergic neurons and is also generated by the absence of one copy of the gene in the same neuronal cell line. In the fear conditioning test, all genotypes showed more freezing during the context phase after conditioning than during the habituation phase and no difference was observed between genotypes in the context response (Supplementary figure 4D). In the sociability 3-chambers test, all the four groups of mice showed preference for the cage containing the mouse rather than the empty cage (Supplementary figure 4E). No difference was found between the four groups in the total amount of time spent sniffing the cage containing the congener (Supplementary figure 4F). Hence, by contrast to *Dyrk1a^C/C^* mice, *Dyrk1a^C/+^* mice do not present decreased social exploratory behaviour. Dp1Yey mice did not spend significantly more time with the novel mouse compared to the familiar one, indicating no preference for social novelty (Figure 4F). This phenotype was rescued by returning to two copies of *Dyrk1a* in glutamatergic neurons (Figure 4F). *Dyrk1a^C/+^* mice also showed preference for social novelty (Figure 4F).

Hence, both increase in *Dyrk1a* copy number in glutamatergic neurons of trisomic mice and haploinsufficiency of *Dyrk1a* in glutamatergic neurons impact explicit memory supporting a key role of *Dyrk1a* in glutamatergic function as a modulator of explicit memory, but other functions probably require normalization of *Dyrk1a* in other cell types to be restored.

### Proteomic analysis confirms the impact of *Dyrk1a* gene dosage on synaptic activity

To examine the contribution of DYRK1A in molecular pathways linked to the cognitive phenotypes associated to T21 in the glutamatergic neurons, we performed proteomic profiling of the hippocampus of control, Dp1Yey, Dp1Yey/*Dyrk1a^C/+^* and *Dyrk1a^C/+^* mice. We identified 63 proteins that were up-regulated and 16 that were down-regulated in the hippocampi of Dp1Yey mice compared with controls. Among those, 40 of the up-regulated and 12 of the down-regulated proteins were back to control levels in Dp1Yey/*Dyrk1a^C/+^* hippocampi, while one up-regulated protein in Dp1Yey was down-regulated in Dp1Yey/*Dyrk1a^C/+^* and 4 down-regulated proteins in Dp1Yey were up-regulated in Dp1Yey/*Dyrk1a^C/+^* mice. We found 51 up-regulated and 7 down-regulated proteins in the hippocampus of *Dyrk1a^C/+^* mice. Eleven of those proteins (CAMK2A, ATP6V1C1, DPP3, ERGIC1, GPM6A, CENPV, RPS28, AGAP2, SNX6, ABCA1, BRK1) were also deregulated in Dp1Yey and back to normal level in Dp1Yey/*Dyrk1a^C/+^*, suggesting that they are impacted by *Dyrk1a* copy number in glutamatergic neurons (Figure 5A and Supplementary Table S4). We performed GO enrichment analysis on the list of deregulated proteins using the ToppCluster website, selecting a Bonferroni cut-off of P<0.05. Enrichment analysis indicates that pathways and GO components that are mostly affected by *Dyrk1a* gene dosage are synaptic, dendritic and axonal components (Figure 5B-C; Supplementary Table S5). Normalization of *Dyrk1a* copy number in the glutamatergic neurons did not rescue specific pathways but had a more global effect with 50 to 80% of the proteins present in each Dp1Yey enriched GO returning to normal amount in Dp1Yey/*Dyrk1a^C/+^* mice (Figure 5B). Interestingly, decreased *Dyrk1a* gene dosage was found to impact pre-synaptic proteins as observed in the transcriptome of *Dyrk1a^C/C^* hippocampi, whereas increased *Dyrk1a* gene dosage was associated with the post-synapse and growth cone (Figure 5C). Proteins enriched in the hippocampus of *Dyrk1a^C/+^* mice were linked to translational activity whereas increased *Dyrk1a* gene dosage was associated to ATPase activity (Figure 5C).

**Figure 5:**
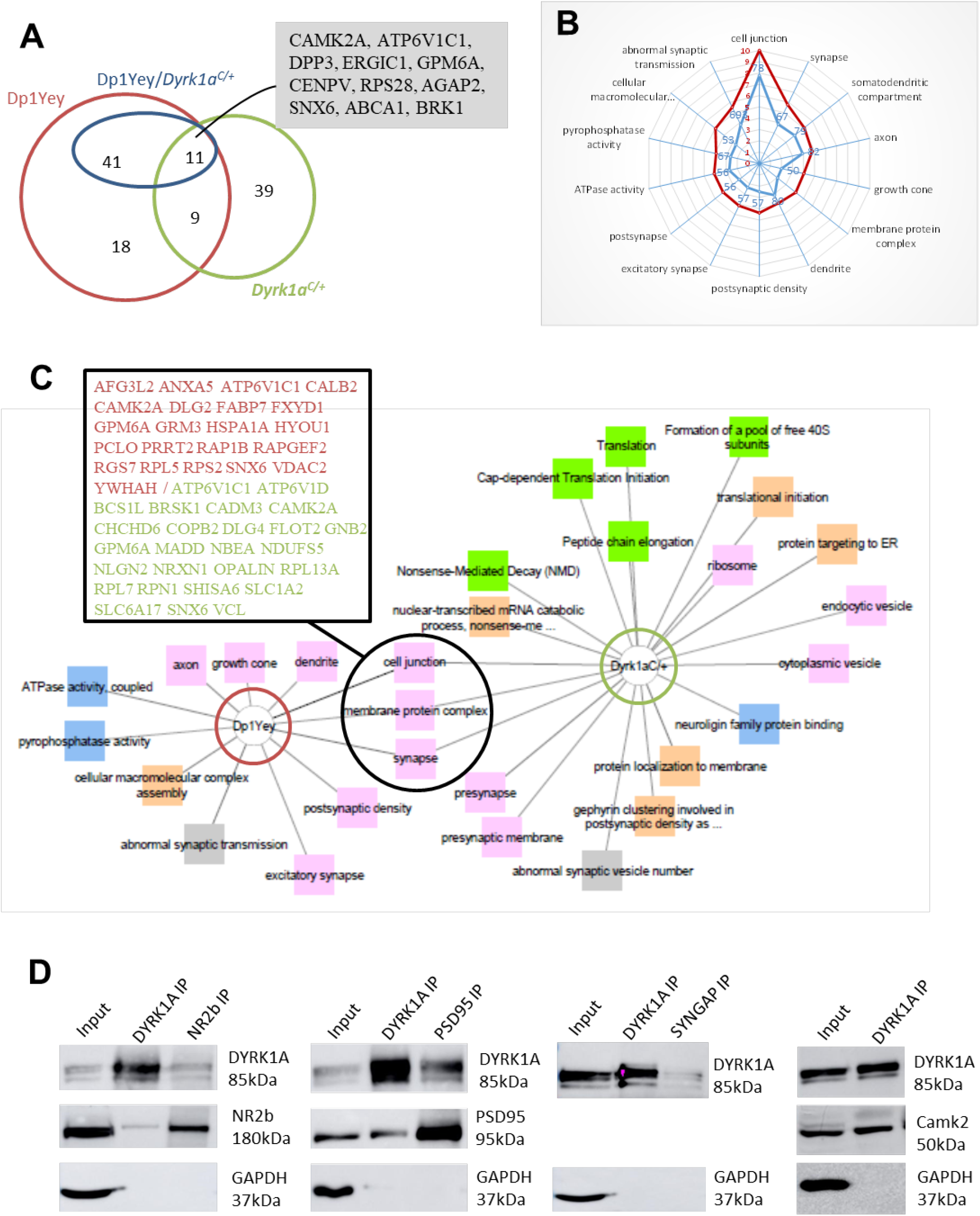
Proteomic analysis. **(A)** Venn diagram showing the numbers of deregulated proteins in the different mouse models. The numbers of proteins shown for the Dp1Yey/*Dyrk1a^C/+^* model (in dark blue) correspond to proteins that were deregulated in the Dp1Yey and back to normal levels in this model (proteins that are regulated by Dyrk1a in the trisomy). Proteins deregulated by both *Dyrk1a* up and down-regulation (common to Dp1Yey, Dp1Yey/*Dyrk1a^C/+^* and *Dyrk1a^C/+^*) are listed in the grey shaded box. (B) Radar plots of GO terms that are mostly enriched in the Dp1Yey model (in red with scale bar corresponding to the log of the p-value) and of the proportion of the proteins found deregulated in Dp1Yey which amount is normalized by the return in 2 copies in the Dp1Yey/*Dyrk1a^C/+^* model (in blue with scale bar corresponding to the % of Dp1Yey deregulated protein back to normal levels). (C) Visual representation of the GO enrichments for the deregulated proteins in Dp1Yey and *Dyrk1a^C/+^* hippocampi with connection between common terms. The different categories of GO are represented by different colors: pink for Cellular components”, blue for “Molecular functions”, green for “Pathways”, orange for “Biological processes” and grey for “Phenotypes”. The list of proteins in the box corresponds to proteins present in the common deregulated GO terms (in red deregulated proteins in Dp1Yey and in green proteins deregulated in *Dyrk1a^C/+^*). (D) Western blots of DYRK1A, NR2B, PSD95, SYNGAP and CAMK2 proteins following IPs of wild-type mice brain extracts. We found in NR2B, PSD95 and CAMK2 in the IPs of DYRK1A. We also detected DYRK1A in the IPs of NR2B, PSD95 and SYNGAP.

### Interaction of DYRK1A with post-synaptic proteins

Behavioural and proteomic analyses suggest a direct impact of DYRK1A at the glutamatergic synapse. Previous work from our laboratory already demonstrated a role of DYRK1A at the presynapse by showing interaction of DYRK1A with SYN1, a neuronal phosphoprotein associating with the cytoplasmic surface of the presynaptic vesicles and tethering them to the actin cytoskeleton [29, 30], and with CAMK2 that was previously shown to phosphorylate SYN1 leading to the release of the vesicle pool [31–33]. Moreover, we also found that SYN1 was phosphorylated by DYRK1A on its S551 residue *in vitro* and *in vivo*, highlighting the role of DYRK1A in SYN1-dependent presynaptic vesicle trafficking [33]. CAMK2, deregulated in our proteomic analysis, is also present in the glutamatergic postsynapse and has a major role in the molecular cascade leading to LTP [34]. To investigate a potential role of DYRK1A at the postsynapse, we looked at DYRK1A protein interaction with CAMK2 and key proteins of the postsynaptic density complex (PSD), GLUN2B (NR2B), PSD95 and SYNGAP. We carried co-immunoprecipitation (co-IP) experiments with adult mouse brain lysates using antibodies against DYRK1A and these proteins and using GAPDH as a negative control. We found CAMK2A, NR2B and PSD95 present in the immunoprecipitates (IPs) of DYRK1A, while DYRK1A was found in the IPs of NR2B, PSD95 and SYNGAP (Figure 5D), showing that these proteins interact together.

## Discussion

Complete *Dyrk1a* inactivation leads to early embryonic lethality with homozygous null *Dyrk1a* mice presenting drastic developmental growth delay with smaller brain vesicles, hindering the investigation of *Dyrk1a* function in the brain [19]. We therefore used a conditional knockout strategy to analyse *Dyrk1a* function in glutamatergic neurons. We found a significant reduction of about 10% of brain weight and size of *Dyrk1a^C/C^* mice compared to control animals. In comparison with *Dyrk1a^+/−^* mice that have 30% brain reduction [35] and consistent with postnatal expression of *Camk2a*, this suggest that microcephaly observed in MRD7 results from different impacts of DYRK1A on brain neurogenesis during embryonic and postnatal development, with *Dyrk1a^C/C^* brain revealing the impact of DYRK1A on postnatal neuronal morphogenesis. Indeed, the cortical size reduction that we observed was associated with increased cell density, as observed by Guedj and collaborators in *Dyrk1a^+/−^* mice [35] and suggesting a reduction of neuronal processes as observed in the neocortex of *Dyrk1a^+/−^* mice [12, 36, 37].

DYRK1A deficit in glutamatergic neurons resulted in specific impacts on mouse behaviour, learning and memory. *Dyrk1a^C/C^* mice were less anxious, spending more time in the centre of the OF and in the open arms of the EPM, and showed decreased freezing performance in the fear-related contextual test indicating an impact on emotional behaviour. Deficit in contextual fear behaviour might be attributed to a defective hippocampal-to-basolateral amygdala transmission as a result of either a deficit in glutamatergic projections or deficit in excitatory activity [38]. Change in emotional behaviour in *Dyrk1a^C/C^* mice is not the result of an intrinsic hyperactivity, as the mice did not present increased locomotor activity either spontaneous (circadian activity) or novelty induced (OF). Hence, contrary to the hypoactivity induced by full *Dyrk1a* haploinsufficiency [39], absence of DYRK1A in glutamatergic neurons does not impact mouse locomotor activity. Thermal pain sensitivity was altered in *Dyrk1a^C/C^* mice that were more sensitive to heat. This higher nociception response was also observed in *Dyrk1a^+/−^* mice, suggesting that DYRK1A has an impact on central processes involved in the control of pain sensitivity. The glutamatergic system takes part in the nociceptive circuits and activation of the expression of IEGs, whose expression was found increased in *Dyrk1a^C/C^* mice, has been shown to be part of long-term events triggered in neuroadaptation to pain in those circuits [40]. Interestingly, we and our collaborators observed a decreased in the expression of some of these IEGs (*Npas4, Arc, c-Fos* and *Fosb*) in the hippocampi of Tg(Dyrk1a) and of the trisomic mouse models Dp1Rhr and Ts65Dn [41] (Duchon et al). IEGs are also believed to be crucial in the formation of long-term memory which we also found impacted in *Dyrk1a^C/C^* and trisomic mice [42]. Moreover, our meta-analysis of the transcriptomic data of hippocampi from five DS mouse models carrying Mmu16 segmental duplications and a transgenic model overexpressing *Dyrk1a* revealed regulatory protein networks centred around six protein hubs, among which were DYRK1A itself and NPAS4 (Duchon et al.) *Npas4* is a neuron-specific gene and is present in both excitatory and inhibitory neurons, activating distinct programs of late-response genes promoting inhibition onto excitatory neurons and excitation on inhibitory neurons [27]. But we did not found major changes in the expression of late response genes targeted by NPAS4 (Supplementary Table S3) [27]. This could be due to experimental bias as the transcriptomic analysis was done using RNA extracts from whole hippocampi containing different cell populations. This heterogeneity can hinder glutamatergic-specific expression of the late-response genes. Even though IEGs are well known markers to measure neuronal activity during cognitive stimulation, their impact in cognitive processes affected in cognitive deficit disorders are unknown and we have no explanation for IEGs overexpression in the hippocampus of *Dyrk1a^C/C^* animals. In our analysis, IEG expression changes have been observed in “naïve” mice that were not subjected to any exercise or behavioural test. It would be therefore also interesting to analyse the expression of IEGs and late-response genes in the mice after induction of neuronal activity, as it was explored for activation of Arc mRNA transcription in pyramidal neurons of the CA1 region of the hippocampus in Ts65Dn mice [43]. In addition to transcriptional up-regulation, we also observed a down-regulation of genes for proteins involved in neuron projection, reinforcing the conclusion of a defective synaptogenesis in *Dyrk1a^C/C^* animals, and proteins involved in synaptic vesicle cycle, implicating DYRK1A in neurotransmitter release. Among the deregulated genes, we found *Amphiphysin* (*Amph*), which its protein is a known target of DYRK1A [44], and *Synapsin 2* (*Syn2*), which paralog SYNAPSIN 1 was found to be hyperphosphorylated by DYRK1A overexpression in TgDyrk1a mice [33]. Moreover, DYRK1A was found to phosphorylate MUNC18-1 [45], which interacts with the SNARE complex protein Syntaxin 1A, whose transcripts are also decreased in the hippocampus of *Dyrk1a^C/C^* mice and which was found to be one of the six hubs connecting the major subnetwork biological cascades found deregulated in DS models (Duchon et al.). Our results together with others [44, 46, 47] point at a role of DYRK1A in the glutamatergic presynapse in the control of neurotransmitter release through synaptic vesicles exocytosis and vesicles recycling processes.

We assessed the impact of *Dyrk1a* deficit in the glutamatergic neurons in *Dyrk1a^C/+^* as a model of MRD7. While *Dyrk1a^C/C^* mice were less anxious, this phenotype was not observed in *Dyrk1a^C/+^* mice, indicating that haploinsufficiency of *Dyrk1a* is not sufficient to trigger this decreased anxiety pattern. Working memory was not affected neither in *Dyrk1a^C/+^* nor *Dyrk1a^C/C^* mice. This fits with our observation that working memory is also not affected in *Dyrk1a^+/−^* mice. We found that long-term recognition memory is affected in both *Dyrk1a^C/+^* and *Dyrk1a^C/C^* mice. Deficit in long-term NOR has also been described in *Dyrk1a^+/−^* mice [12] and in transgenic model overexpressing *Dyrk1a* alone [33] (DUCHON). Together with the rescue of this type of memory in Dp1Yey, our findings show that DYRK1A has a direct cell-autonomous function in regulating long-term explicit memory in glutamatergic neurons.

Cognitive deficits observed in DS have been linked to a perturbation of synaptic transmission due to defects in the control of the excitatory/inhibitory balance. In addition, most of the phenotypes observed in DS people and models are due to defects in the hippocampus or the prefrontal cortex [48]. Analysis of trisomic mouse models has revealed an overproduction of the inhibitory neurotransmitter GABA restricting synaptic activation of the glutamatergic NMDA receptors. Furthermore, *Dyrk1a* has been implicated in glutamate-GABA imbalance [49, 50]. However, how *Dyrk1a* controls the balance between the two pathways is still not clear. We performed a genetic rescue returning to two copies of *Dyrk1a* exclusively in cortical and hippocampal glutamatergic neurons of Dp1Yey mice and carried behavioural analysis of those mice to see if this rescue was enough to restore cognitive functions in Dp1Yey mice. We already found that Dp1Yey mice have deficits in both hippocampus-dependent spatial working and long-term explicit memory (Duchon et al.). In addition, Yu and collaborators showed deficits in hippocampal-mediated context memory [51]. We confirmed the deficit in in long-term explicit memory and in working memory. However, we did not observe a deficit in contextual memory in those mice. Normalization of gene copy number in glutamatergic neurons only partly restored the cognitive functions that were impacted in the Dp1Yey mice. Working memory deficit was not restored in Dp1Yey/*Dyrk1a^C/+^* mice. As *Dyrk1a* haploinsufficiency does not impact working memory, this could question the role of *Dyrk1a* in the deficit observed in Dp1Yey mice. Nevertheless, the overexpression of *Dyrk1a* alone was able to induce a deficit in spontaneous alternation [33] (DUCHON). Deletion of *Dyrk1a* under Camk2a-Cre occurs after birth and hence the lack of rescue of working memory in Dp1Yey/*Dyrk1a^C/+^* mice could result from prenatal brain defects. However, it was shown that spontaneous alternation in rodents is not present during the early postnatal stages of development (before postnatal day 30), indicating that brain processes sustaining this behaviour develops between the second and fourth postnatal week [52]. Furthermore, treatment of adult mice with an inhibitor of DYRK1A is sufficient to restore working memory in Ts65Dn mice [50]. This lack of rescue presumably comes from the effect of *Dyrk1a* overexpression in non-glutamatergic neurons. The role of the GABAergic system on spontaneous alternation is not determined but the injection of a GABA_A_ receptor agonist has been shown to decrease spontaneous alternation rates [53, 54]. Furthermore, DYRK1A was shown to act on GABA-producing enzymes [49]. Testing this hypothesis will require to normalize *Dyrk1a* in GABAergic neurons.

DS patients are affected in their explicit long-term memory abilities with a particular impairment in the visuo-perceptual processing [55]. We used the visual-object recognition NOR test as a paradigm to assess long-term explicit memory in the mouse. Both TgDyrk1a and Ts65Dn mouse models have been shown to have impaired long-term object recognition memory that could be ameliorated by treating the adult mice with the DYRK1A inhibitors [56–58]. We show here that correcting *Dyrk1a* gene copy number in glutamatergic neurons is sufficient to rescue explicit long-term memory in Dp1Yey/*Dyrk1a^C/+^* mice. Moreover, DYRK1A shortage in glutamatergic neurons is also sufficient to trigger long-term memory deficit. Hence *Dyrk1a* gene dosage seems to have an important role in glutamatergic neuronal defects observed in DS and MRD7 mouse models. TgDyrk1a mice have been shown to have bidirectional changes in synaptic strength with elevated LTP, reduced LTD [10] and dysregulated NMDA-receptor mediated calcium signalling [59]. Furthermore, normalization of *Dyrk1a* expression in the hippocampus of Ts65Dn mice can partially restore the deficit of LTP in the CA1 of Ts65Dn mice. As Dp1Yey mice show similar hippocampal LTP deficit [51], it would be interesting to see if *Dyrk1a* normalisation in the glutamatergic neurons could restore LTP in Dp1Yey/*Dyrk1a^C/+^* mice.

Excessive GABAergic inhibition has been proposed as the major cause of the perturbation between excitatory and inhibitory neurotransmission, with glutamatergic deficit being the consequence of over-inhibition of the NMDA receptors resulting in deficit of LTP and memory [60]. Our finding outlines the glutamatergic deficit as a distinct alteration with *Dyrk1a* overexpression playing a key role in glutamatergic dysfunction and GABA-mediated over-inhibition combining with it to produce the full DS cognitive deficit. This also raises the question of the role of *Dyrk1a* overexpression in GABAergic neurons as other trisomic genes are also potential candidates for neuronal dysfunction. For example, overexpression of *Girk2* leads to increase in GABA_A_-mediated GIRK currents in hippocampal neuronal cultures, affecting the balance between excitatory and inhibitory transmission [61, 62].

The finding of a cell-autonomous impact of DYRK1A in glutamatergic neurons on long-term memory function is supported by the impact of increased *Dyrk1a* gene dosage in glutamatergic neurons on the amount of glutamatergic post-synaptic proteins. Hence, among enriched proteins in the hippocampus of Dp1Yey mice that turned back to normal in the hippocampus of Dp1Yey/*Dyrk1a^C/+^* mice, we found CAMK2A, a subunit of the calcium/calmodulin-dependent protein kinase II (CAMK2) which plays a critical role in LTP by regulating ionotropic glutamate receptors at postsynaptic densities, GPM6A, a neuronal membrane glycoprotein involved in neuronal plasticity, regulation of endocytosis and intracellular trafficking of G-protein-coupled receptors [63], the GRM3 G-protein-coupled metabotropic glutamate receptor, DLG2, a member of the postsynaptic protein scaffold of excitatory synapses interacting with the cytoplasmic tail of NMDA receptors [64] and the intracellular calcium-binding protein CALB2 functioning as a modulator of neuronal excitability [65]. Previous work done in our laboratory found several proteins from the PSD that were hyperphosphorylated in mice with three copies of *Dyrk1a* (TgDyrk1a) [35] and dephosphorylated by TgDyrk1a mouse treatment with a DYRK1A inhibitor [33]. Among those proteins, the NR2B is a subunit of the glutamatergic postsynaptic NMDA receptor which play a pivotal role in excitatory synaptic transmission. This result was validated by our Co-IP experiments showing an interaction between DYRK1A and NR2B. NR2B subunits are expressed in the neocortex and hippocampus [66–68]. NMDA receptors in the mature hippocampus consist of two NR1 subunits associated with either two NR2A, two NR2B or one of each subunits [69, 70] and different forms of synaptic plasticity have been associated to different types of NMDA receptors [71–73]. Hence, in addition to its interaction with the NR1/NR2A-type of receptors [74], we also point out an association with NR1/NR2B receptors. Moreover, interaction between DYRK1A and the PSD proteins PSD95, CAMK2 and SYNGAP, detected by coIP, strongly suggests a role of DYRK1A at the glutamatergic post-synapse. In addition, absence of the GLUR1 subunit of the AMPA receptor in the IP of DYRK1A indicates that DYRK1A interact most specifically with the NMDA-PSD complex. Altogether, this strongly suggests an implication of DYRK1A at the glutamatergic post-synapse, somehow supporting its involvement in long-term memory formation.

Taking advantage of a conditional allele for *Dyrk1a* inactivation, we were able to associate *Dyrk1a* gene dosage changes in glutamatergic neurons to specific cognitive phenotypes and molecular modifications and demonstrated a major impact of *Dyrk1a* dose change at the glutamatergic synapse on long-term explicit memory while no impact was observed for motor activity, short-term working memory and susceptibility to epilepsy. Further analysis of DYRK1A impact on other neurons, such as GABAergic ones, will be necessary to understand how DYRK1A perturbs the excitatory/inhibitory pathways, resulting in the full DS and MRD7 cognitive deficits.

## Materials and Methods

### Mouse lines

The Dp(16Lipi-Zbtb21)1Yey (Dp1Yey) line was created by Yu and collaborators [75] and bears a 22.6 Mb segmental duplication of the *Lipi-Zfp295* fragment of murine chromosome 16 syntenic to Hsa21 [51, 75]. The transgenic Tg(Camk2-Cre)4Gsc line [20, 76] expressing the Cre recombinase under the control of the Camk2a promoter to inactivate the targeted conditional knockout allele in glutamatergic neurons of the cortex and hippocampus after birth. The Dyrk1^tm1.ICS^ conditional knockout (noted Dyrk1a cKO) was generated at the PHENOMIN-ICS (Institut Clinique de la Souris; Illkirch, France; www.phenomin.fr) in the frame of the Gencodys consortium (http://www.gencodys.eu/).The targeting vector was constructed as follows. A 1096 bps fragment encompassing exon 7 (ENSMUSE00001246185) was amplified by PCR (from BAC RP23-115D20 genomic DNA) and subcloned in an MCI proprietary vector. This MCI vector contains a LoxP site as well as a floxed and flipped Neomycin resistance cassette. A 3.8 kb fragment corresponding to the 3’ homology arm and 4.1 kb fragment corresponding to the 5’ homology arms were amplified by PCR and subcloned in step1 plasmid to generate the final targeting construct. The linearized construct was electroporated in C57BL/6N (B6N) mouse embryonic stem (ES) cells. After selection, targeted clones were identified by PCR using external primers and further confirmed by Southern blot with a Neo probe (5’ and 3’ digests) as well as a 5’ external probe. Two positive ES clones were injected into BALB/cN blastocysts. Resulting male chimeras were bred with Flp deleter females previously backcrossed in a C57BL/6N [77] (PMID: 10835623). Germline transmission of the conditional allele was obtained (Figure 1B). The Flp transgene was segregated by a further breeding step. By combining the three different lines together, we obtained the following groups of mice for phenotyping analyses: Dp1Yey (trisomic, for the *Lipi-Zfp295* fragment containing the *Dyrk1a* gene), Dp1Yey/*Dyrk1a^C/+^* (trisomic for the *Lipi-Zfp295* fragment but containing only two copies of *Dyrk1a* in glutamatergic neurons), *Dyrk1a^C/+^* (containing only one copy of *Dyrk1a* in the glutamatergic neurons) and *Dyrk1a^C/C^* (knocked out for *Dyrk1a* in glutamatergic neurons). Wild-type, *Dyrk1a^cKO/+^* and *Dyrk1a^cKO/cKO^* mice were used as disomic controls.

For the genotyping of the mice and identification of the *Dyrk1a* knockout allele in the brain, genomic DNA was isolated from tail and different organ biopsies using the NaCl precipitation technique. 50-100 ng of genomic DNA was used for PCR. Primers used for the identification of each allele and size of PCR products are described in Figure 1 and Supplementary Table S6. Details on the genotyping protocol used here are published [78].

The mice were housed in groups (2–4 per cage) and were maintained under specific pathogen-free (SPF) conditions and were treated in compliance with the animal welfare policies of the French Ministry of Agriculture 133 (law 87 848).

### Mouse RT droplet digital PCR (ddPCR)

Total RNA was extracted from frozen brain tissues (cerebellum, cortex, striatum, hippocampus and thalamus/hypothalamus) of five wt and four *Dyrk1a^C/C^* mice as described in Lindner *et al*. 2020 [79]. For ddPCR, all primers were designed and synthesized as described in Lindner *et al*. 2020 [79] excepting Universal Probe Library probe used to *Dyrk1a* mRNA which is provided by Roche. *Dyrk1a* and *Hprt* primers and probes sequences are given in Supplementary Table S6. RNA reverse transcription, droplet generation, PCR amplification, droplets quantification and analysis are also described in Lindner *et al* 2020 [79]. We presented the results as a ratio of the mean of *Dyrk1a* RNA transcript in *Dyrk1a^C/C^* tissue normalized to the mean of *Dyrk1a* RNA transcript in wt tissue. Experiments were performed following dMIQE guidelines for reporting ddPCR experiments (Supplementary Table S7) [79, 80].

### Western blot analysis

Twenty-five microgram of total protein extracts from hippocampi (n=3 per genotype) were electrophoretically separated in SDS-polyacrylamide gels (10%) and transferred to a nitrocellulose membrane (100V, 2h at room temperature). Non-specific binding sites were blocked with 5% skimmed milk in Tween20 0.1% Tris buffer saline 1h at room temperature. Immunostaining was carried out with a mouse monoclonal anti-Dyrk1a (Abnova, H00001859-M01) and an anti-Gapdh antibodies (ThermoFisher, MA5-15738), followed by secondary anti-mouse IgG conjugated with horseradish peroxidase (DAKO). The immunoreactions were visualized by ECL chemiluminescence system (Amersham) with the Amersham Imager 600. Semi-quantitative analysis was performed using ImageJ software (W. Rasband, NIH; http://rsb.info.nih.gov/ij/).

### Immunohistological analysis

Adult mice were deeply anesthetized with sodium pentobarbital and perfused intracardially with 30 ml PBS followed by 30 ml 4% paraformaldehyde in PBS. Brains were removed from the skull and immersed in the same fixative overnight. After rinsing with PBS, the brains were transferred into 70% ethanol until paraffin inclusion. For inclusion, brains were dehydrated and embedded in paraffin. Serial 10 μm sections were made with a microtome.

Brain sections were stained using the myelin-specific dye luxol fast blue and the Nissl staining cresyl violet. Briefly, brain sections were deparaffinised, rehydrated and incubated in 0.1% luxol fast blue (95% alcohol and 0.5% acetic acid) solution at room temperature overnight. After rinsing excess stain with 95% ethanol and deionized water, the slides were placed in 0.05% lithium carbonate solution for 10 seconds followed by 70% ethanol for 5 seconds. They were then rinsed in deionized water until the colourless grey matter contrasted with the blue-green white matter. Sections were then stained in 0.1% cresyl violet acetate solution for 5 minutes at 56°C in a water bath, rinsed in deionized water and quickly in 100% ethanol. Sections were dried, cleared in histosol^®^ (Shandon) and mounted in Eukitt^®^ (Labonord).

Immunohistology was performed using a standard protocol. After deparaffinization, rehydration and antigen retrieval (10 mM citric acid, 0.05% Tween 20, pH6.0) for 45 min in a 94°C water bath, sections were incubated in a blocking solution (0.05% Tween20, 5% Horse serum) for 1 hour at room temperature. Sections were then incubated at 4°C overnight with the primary antibodies (Mouse anti-Dyrk1a: Abnova, Cat. N. H00001859-M01; Rabbit anti-Camk2a: Molecular probes, PA5-14315). After washing, sections were incubated with anti-Mouse and anti-Rabbit Alexa Fluor^®^546 or 488 secondary antibodies for detection. The sections were mounted with Mowiol mounting medium (0.1M Tris (pH8.5, 25% glycerol, 10% w/v Mowiol 4-88 (Citifluor)) containing DAPI (5 μg/ml) and images were acquired using Hamamatsu Nanozoomer 2.0 (Hamamatsu, Hamamatsu City, Japan) and a Leica Upright fluorescent microscope (Leica Microsystems, Heidelberg).

Immunohistochemistry was performed using a standard protocol. Briefly, antigen retrieval was performed by heating the slides in Tris/EDTA buffer (10 mM Tris Base, 1 mM EDTA, 0.05% Tween 20, pH 9.0) for 45 min in a 94°C water bath. Then, the sections were quenched in 0.3% oxygen peroxide solution for 20 min and blocked with 10% normal horse serum and 0.1% Triton X-100 in 1× PBS for 1 h at room temperature. The sections were incubated overnight at 4°C with a rabbit anti-Olig2 antibody (1:500, Santa Cruz sc-48817). which was detected by incubating the sections with secondary biotinylated antibodies (Life Technologies™) for 2 h at room temperature and then with an avidin-biotin complex at 37°C for 30 min. Dark coloration was developed with diaminobenzidine tetrahydrochloride and the sections were mounted with aqueous mounting medium (Faramount aqueous mounting medium, Dako^®^).

### Morphometric analysis

Morphometric analysis was performed on three *Dyrk1a^C/C^* and three control mice based on the standard operating procedures for morphological phenotyping of the mouse brain using basic histology [81]. Surface and cortical thickness measurements as well as cell counting were conducted on scanned images using Hamamatsu Nanozoomer 2.0 from luxol fast blue/cresyl violet-stained sections around Bregma −1.5 mm (Paxinos adult mouse brain atlas, Franklin and Paxinos, 1997). TIFF files were opened in ImageJ with the following settings: 9 decimal places (using the panel Analyze/Set Measurements) and “cm” as unit length (using Analyze/Set Scale). The polygone selection tool was used to measure area and the straight line tool was selected to measure length. The thickness of the different cortical layers (layer I to layer VI) were estimated in the somatosensory cortex based on the shape and density of the neurons on these different layers. Cell count performed in the somatosensory cortex was done within a counting frame of 0.1 cm. Cell count performed in the CA1 was done within a counting frame of 0.04 cm width). Olig2+-positive cells within the corpus callosum were counted by measuring a distance of 1 mm from the midline of the brain and selecting the corpus callosum area underneath. Cell count was done manually.

### RNA-seq libraries and analysis

Total RNA was Trizol-extracted from 2 wild-type and 2 Dyrk1a^*C/C*^ frozen P30 hippocampi. RNA was treated with DNase (Qiagen) and purified on the RNeasy MinElute Cleanup Kit (Qiagen). 2 μg of total RNA were treated with the Ribo-Zero rRNA Removal Kit (Human/Mouse/Rat; Illumina). Depleted RNA was precipitated 1h at −80°C in three volumes of ethanol plus 1 μg of glycogen. RNA was then washed and resuspended in 36 μl of RNAse free water. RNA fragmentation buffer (NEBNext^®^ Magnesium RNA Fragmentation Module) was added to the solution and the RNA was fragmented by incubation at 95°C for 3 min. cDNA first strand synthesis was performed with random hexamer primers and cDNA second strand synthesis was performed with dUTPs, to ensure strand specificity. The RNA-seq library was synthetized with KAPA Hyper prep kit (Kapa Biosystems, Wilmington, MA, USA): a treatment with USER enzyme (NEB, M5505L) was added to digest the unspecific strand.

The libraries were pooled (4/lane) on an Illumina HiSeq. 2000. Libraries were sequenced (50 cycles, single-end) yielding on average 40 million mapped reads. RNA-Seq libraries were mapped with GSNAP (version 2015-06-23) against mm9 mouse RefSeq annotations updated to the 28/7/2015.

DESeq 2 (v1.14) was used to perform statistical comparisons. All the enrichment analysis were made from standard hypergeometric tests with benjamini or bonferroni correction. The markers of hippocampal cell types were obtained from [ref] and the common background genes were evaluated prior to the enrichment (hypergeometric test). GO annotations were updated to 25/6/2015.

### Proteomic analysis

Fourty microgram of total protein extracts from hippocampus (4 controls, 5 Dp1Yey, 4 Dp1Yey; *Dyrk1a^C/+^* and 5 *Dyrk1a^C/+^*) were used for the preparation. Samples were precipitated, reduced, alkylated and digested with LysC and trypsin at 37°C overnight. 10 μg of each sample were then labeled with TMT isobaric tags, pooled, desalted on a C18 spin-column and dried on a speed-vacuum before nanoLC-MS/MS analysis. Samples were separated on a C18 Accucore nano-column (75 μm ID x50 cm, 2.6 μm, 150 Å, Thermo Fisher Scientific) coupled in line with an Orbitrap ELITE mass spectrometer (Thermo Scientific, San Jose, California). Samples were analyzed in a Top15 HCD (High Collision Dissociation) mass spectrometry on 8h gradient. Data were processed by database searching using SequestHT (Thermo Fisher Scientific) with Proteome Discoverer 1.4 software (Thermo Fisher Scientific) against a mouse Swissprot database (release 2015-03). Peptides were filtered at 5% false discovery rate (FDR) and one peptide in rank 1. Protein quantitation (ratio of the intensity of the fragmented tag in sample “x” to the intensity of the fragmented tag in one control (disomic) sample used as the reference) was performed with reporter ions quantifier node in Proteome Discoverer 1.4 software with integration tolerance of 20 ppm, and the purity correction factor were applied according to the manufacturer’s instructions. A scaling factor normalization method was used in order to make sample ratios comparable. Ratios were normalized by calculating the mean of all the peptide ratios in one sample, calculating a scaling factor (sf=mean [ratio control ref]/mean [ratio sample x]) for each sample and multiplying each ratio by the sf. Data were filtered with the following criteria: minimum number of peptide ratios used to calculate the protein ratio equal to 2; variability of the peptide ratios <20%; ratio of Dp1Yey and *Dyrk1a^C/+^* samples compared to mean of disomic controls, x>1.2 or x<0.8 and ratio of Dp1Yey;*Dyrk1a^C/+^* samples compared to mean of disomic controls 0.8<x<1.2 among proteins that were selected as deregulated in Dp1Yey. GO enrichment was calculated in the ToppCluster website (https://toppcluster.cchmc.org/), looking at enrichment within the following features: Molecular functions, Biological processes, Cellular components, Phenotypes and Pathways, and using a Bonferroni correction cut-off of P<0.05. The results of the enrichments can be found in the Supplementary Table, Excel File S5.

### Co-immunoprecipitation

Immunoprecipitations were performed on fresh half brains of 3-month old wild-type male mice. Brains were dissected and lysed in 1.2 ml RIPA lysis buffer (Santa-Cruz Biotechnology, France) using Precellys^®^ homogenizer tubes. After centrifugation at 2800 g for 2×15 s, 1 ml brain extract was incubated with 2 μg of antibody of interest at 4°C for 1 h under gentle rotation. An aliquot of the remaining supernatant was kept for further immunoblotting as homogenate control. Then, 20 μl protein G agarose beads, previously washed three times with bead buffer, were added to the mix and gently rotated at 4°C for 30 min. After a 1 min spin at 10,000 g and removal of the supernatant, the pelleted immune complexes were washed three times with bead buffer before WB analysis with appropriate antibodies directed against DYRK1A (H00001859 M01, Interchim; 1:1000), NMDAR2B (Abcam, #ab65783), PSD95 (ab18258, Abcam, France; 1:1000), CAMK2A (PA5-14315, Thermo Fisher Scientific; 1:1000), SYNGAP (sc-8572, Santa Cruz biotechnologies; 1:5000) and GAPDH (MA5-15738, Thermo Fisher Scientific; 1:3000). Immunoblots were revealed with Clarity Western ECL Substrate (Bio-Rad).

### Mouse behavioural analysis

A series of behavioural experiments were conducted in mice with a range of age starting at 2,5 up to 7 months, as described in the Supplementary information. For all these tests, mice were kept in ventilated cages with free access to food and water. The light cycle was controlled as 12 h light and 12 h dark (lights on at 7:00 AM) and the tests were conducted between 8:00 AM and 4:00 PM. Due to the difficulty to obtain Dp1Yey; *Dyrk1a^KO/+^* mice (triple crossing and subfertility of the Dp1Yey line), both males and females were pooled for the analysis. Animals were transferred to the experimental room 30 min before each experimental test. Behavioural experimenters were blinded as to the genetic status of the animals. Behavioural experiments were performed in agreement with the EC directive 2010/63/UE86/609/CEE and was approved by the local animal care, use and ethic committee of the IGBMC (Com’Eth, no.17, APAFIS 2012-069). The PTZ-induced seizures protocol received the accreditation number APAFIS#6321. All the standard operating procedures for behavioural phenotyping have been already described [82–84] and are detailed in the supplementary information.

### Statistical analysis

Statistical analyses were performed using SigmaPlot software. For histological assessments and behavioral tests comparing *Dyrk1a^Camk2aCre/Camk2aCre^* animals to controls, statistical analyses were performed using unpaired t-test when appropriate or the non-parametric Mann-Whitney rank sum test unless otherwise stated in the text. For the four groups analyses (Dp1Yey, Dp1Yey, Dp1Yey; *Dyrk1a^C/+^*, Dp1Yey, *Dyrk1a^C/+^* and controls) a two-way ANOVA did not reveal a significant effect of sex and no interaction with the genotype. Therefore, the sex factor was dropped from the model and a one-way ANOVA and *post hoc* Tukey’s multiple comparison test were used to analyse differences between the four genotype groups.

## Acknowledgments

We would like to thank members of the research group, of the IGBMC laboratory and of the ICS. We are grateful to the IGBMC proteomic plateform and Doulaye Dembele for their expert technical assistance in proteomic analysis, and Binnaz Yalcin and Stephan Collins for their help in brain morphometric analysis. We extend our thanks to the animal care-takers of the ICS who oversee the mice wellness.

This work has been supported by the National Centre for Scientific Research (CNRS), the French National Institute of Health and Medical Research (INSERM), the University of Strasbourg (Unistra), the French state funds through the “Agence Nationale de la Recherche” under the frame programme Investissements d’Avenir [ANR-10-IDEX-0002-02, ANR-10-LABX-0030-INRT, ANR-10-INBS-07 PHENOMIN to YH]. This project has received funding from the Jérôme Lejeune foundation and the European Union’s Horizon 2020 research and innovation programme under grant agreement No 848077. The funders had no role in study design, data collection and analysis, decision to publish, or preparation of the manuscript.

## Conflict of Interest

SDG is Founder, President and CEO of Proteas Bioanalytics Inc., BioLabs at the Lundquist Institute, 1124 West Carson Street, Torrance, CA 90502, USA.

## Notes

### Competing Interest Statement

One cauthor Spiros D. Garbis is Founder, President and CEO of Proteas Bioanalytics Inc., BioLabs at the Lundquist Institute, 1124 West Carson Street, Torrance, CA 90502, USA.

