## Supplementary information and figures for "*Dyrk1a* gene dosage in glutamatergic neurons has key effects in cognitive deficits observed in mouse models of MRD7 and Down syndrome"

***Corresponding author: Yann Hérault**

### A. Supplementary Materials and Methods

Circadian activity:

Circadian activity: spontaneous locomotor activity and rears were measured using individual boxes equipped with infrared captors. The mice were tested for 35 h to measure apparatus habituation and nocturnal and diurnal activities. The results are expressed per 1-h period.

Open Field:

This test measures exploratory and locomotor activities in a novel environment. The mouse is placed in a 55 cm diameter circular arena for a unique 30 min session during which its activity is recorded with a video tracking system (Ethovision, Noldus, France). Distance travelled and time spent in the different zones of the arena (center and periphery) are recorder. After each mouse trial, the arena was thoroughly cleaned to minimize olfactory cues.

Elevated plus maze:

The apparatus used was completely automated and made of PVC (Imetronic, Pessac, France). It consisted of two open arms (30 X 5 cm) opposite one to the other and crossed by two enclosed arms (30 X 5 X 15 cm) and elevated 66 cm from the floor. The apparatus was equipped with infrared captors allowing the detection of the mouse in the enclosed arms and different areas of the open arms. The light intensity at the extremity of the open arms was kept at 50 Lux. Each mouse was tested for 5 min after being placed in the central platform and allowed to explore freely the apparatus. The number of entries and the time spent in the open arms were used as an index of anxiety. Total arm entries were used as measures of general motor activity. The number of rears in closed arms, as well as ethological parameters such as stretching, attempts and head dips, were also automatically scored.

Rotarod:

This test measures the ability of an animal to maintain balance on a rotating rod (Bioseb, Chaville, France). Mice were given three testing trials during which the rotation speed accelerated from 4 to 40 rpm in 5 min. Trials were separated by 10-15 min interval. The average latency was used as an index of motor coordination performance.

Hot plate test:

The hot plate test is used for evaluation of response to acute thermal pain. The mice are placed into a glass cylinder on a hot plate adjusted to 52°C and the latency of the pain reaction (licking, moving the paws, little leaps) is recorded.

Y-maze:

This test assesses short-term working memory by recording spontaneous alternation in a Y-shaped maze. The maze is made of three arms of 40x9x16 cm set at angles of 120° having different motifs on their walls. The animals are placed in one arm and allowed to freely explore the arms during 5 minutes. The sequence of arm entries (when the mouse is with its four paws in the arm) is recorded and percentage of spontaneous alternation is calculated ([number of sequences of three different arms visited / total of visited arms -2] x100). Total number of arm entries is scored as an index of locomotor activity. Mice that visited less than 10 arms were not included in the spontaneous alternation analysis as a low number of visited arms biases this calculation.

The novel object recognition task:

This is based on the innate tendency of rodents to spend more time exploring novel objects over familiar ones. This test is done in two sessions of 10 min each (starting from the first sniffing of an object) at 24 hours interval. In the first trial, the animals were presented with a pair of identical objects. Animals exploring less than 3 second the two objects during this presentation phase were removed from the test. In the second session (test phase), one of the familiar objects is replaced by a new one, and the animal left to explore. The duration of sniffing either the familiar or the novel object is recorded manually and percentage of time spent on each object is calculated as an index of memory (statistically different form 50% chance). To avoid bias of preference, familiar and new objects are randomly assigned for each mouse as well as their emplacement (left or right). The arena and objects are cleaned between each mouse in order to avoid olfactory bias.

Three-chamber social behaviour test:

This test assesses the social interaction behavior of the mice. The system, available from Stoelting (Dublin, Ireland), is composed of three 300 successive identical chambers (20 cm × 40 × cm 22(height) cm) with 5 cm × 8 cm openings allowing access between the chambers. During the habituation phase, the test mouse is placed in the middle chamber and allowed to explore the three chambers under video tracking for 10 min, with each of the two side chambers containing an empty wire cage. The second phase of the test corresponding to the sociability test is carried out directly after the habituation phase. The test mouse is enclosed in the central box, while an unfamiliar mouse (stranger 1) is placed in one of the wire cages in a random manner. The doors are re-opened and the test mouse is allowed to explore the entire social test box for 10 min. The time spent sniffing each wire cage are recorded. The third phase tests the preference for social novelty. A new stranger mouse (stranger 2) is placed into the empty wire cage and the test mouse is allowed to explore again the entire social test box for 10 min, having the choice between the first, already-investigated mouse (stranger 1) and the novel unfamiliar mouse (stranger 2). The same measures are taken as for the sociability test. For the comparison of Dp1Yey, Dp1Yey/*Dyrk1a*$^{C/+}$ and *Dyrk1a*$^{C/+}$ mice, preference for social novelty was tested during two minutes as no significant difference was obtained for control mice when looking higher exploratory times. The entire social test box is washed between each test mouse to remove odours and the time spent in each lateral chamber during the habituation phase is recorded to check for place preference. Several days before the test, stranger mice are habituated to the test in the wire cage 5 min per day for 5 days. Each stranger mouse is chosen randomly for either the sociability session or the preference for social novelty session.

Fear conditioning:

The fear conditioning is an associative learning paradigm for measuring aversive learning and memory. In the fear conditioning procedure, a neutral conditioned stimulus (CS) such as light or tone is paired with an aversive unconditioned stimulus (US) such as mild foot shock. Concomitantly, animals associate the background context cues with the CS. After conditioning, the CS or the spatial context elicits a central state of fear in the absence of the US, expressed as reduced locomotor activity or total lack of movement (freezing). Immobility time is used as a measure of learning/memory performances. Experiments are conducted in operant dimly lit chambers (28 x 21 x 22 cm) equipped with a metal bar floor linked to a shocker (Coulbourn Instruments, Allentown US), a speaker for tone delivery and an infrared activity monitor. The experimental procedure comprises 3 sessions run over 2 days. In the conditioning session the mouse is allowed to acclimate for 4 min, then a light/tone (10 kHz, 80-dB) CS is presented for 20 s and terminated by a mild (1 s, 0.4 mA) foot shock (US). Animals are left in the cage another 2 minutes. Context testing is performed the next day by placing back the mice in the same environment for 6 min without presentation of the light/auditory CS. The movement of the animal is monitored to detect freezing behavior consequent to recognition of the chamber as the spatial context (contextual learning).

Pentylenetetrazol (PTZ)-induced seizures:

PTZ was dissolved in saline (0.9 % NaCl) and injected intraperitoneally at the dose of 30 mg/kg. Immediately after injection, the mouse was placed into a new cage and observed for at least 20 min. The seizure profile (myoclonic, clonic, tonic) and the latency to clonico-tonic seizure were recorded. Fourty-height hours after the first injection, mice received a second injection of 50 mg/kg and were recorded as described above. Seizures profile was analysed using the qualitative Chi square test.

### B. Supplementary Tables

S1: List of Up regulated genes in *Dyrk1a*$^{C/C}$ hippocampi compared with controls (Deseq algorithm, P<0.025)

S2: List of enriched cell populations in deregulated *Dyrk1a*$^{C/C}$ hippocampal genes

S3: List of early and late response gene deregulated in *Dyrk1a*$^{C/C}$ hippocampus

S4: List of proteins that are impacted by the different genetic conditions in the proteomic analysis

S5: List of GO and pathways enriched in the proteome analysis

S6: List of primers and probes used for genotyping and QRT-PCR analysis

**C. Supplementary Figures**

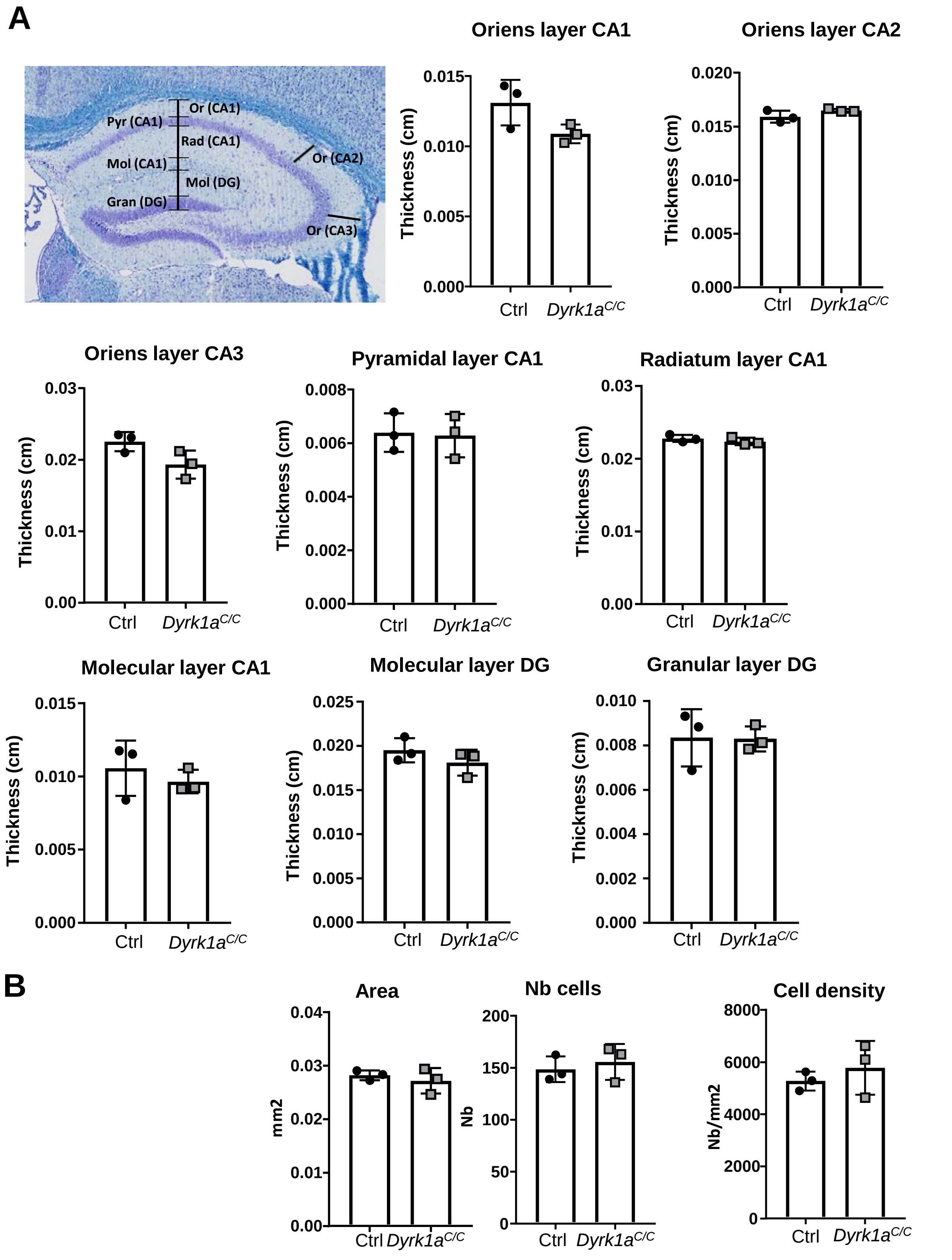


**Supplementary figure 1: Histomorphological analysis of the hippocampus of *Dyrk1a*$^{C/C}$ mice.**

(A) Representative coronal section of hippocampus at Bregma -1.5 stained with cresyl violet and luxol blue that were used for measurements (Magnification 20X) and dot plots indicating the thickness of the different cellular and molecular layers. (B) Enlarged image of the CA1 showing the selected area made for counting the number of cells within the CA1 and dot plots for the area of the CA1, the number of cells within this area and the cell density. Data are presented as point plots with mean ± SD (n=3 females per genotype). Pyr: pyramidal layer, Mol: molecular layer, Gran: granular layer, Or: oriens layer, Rad: radiatum layer.

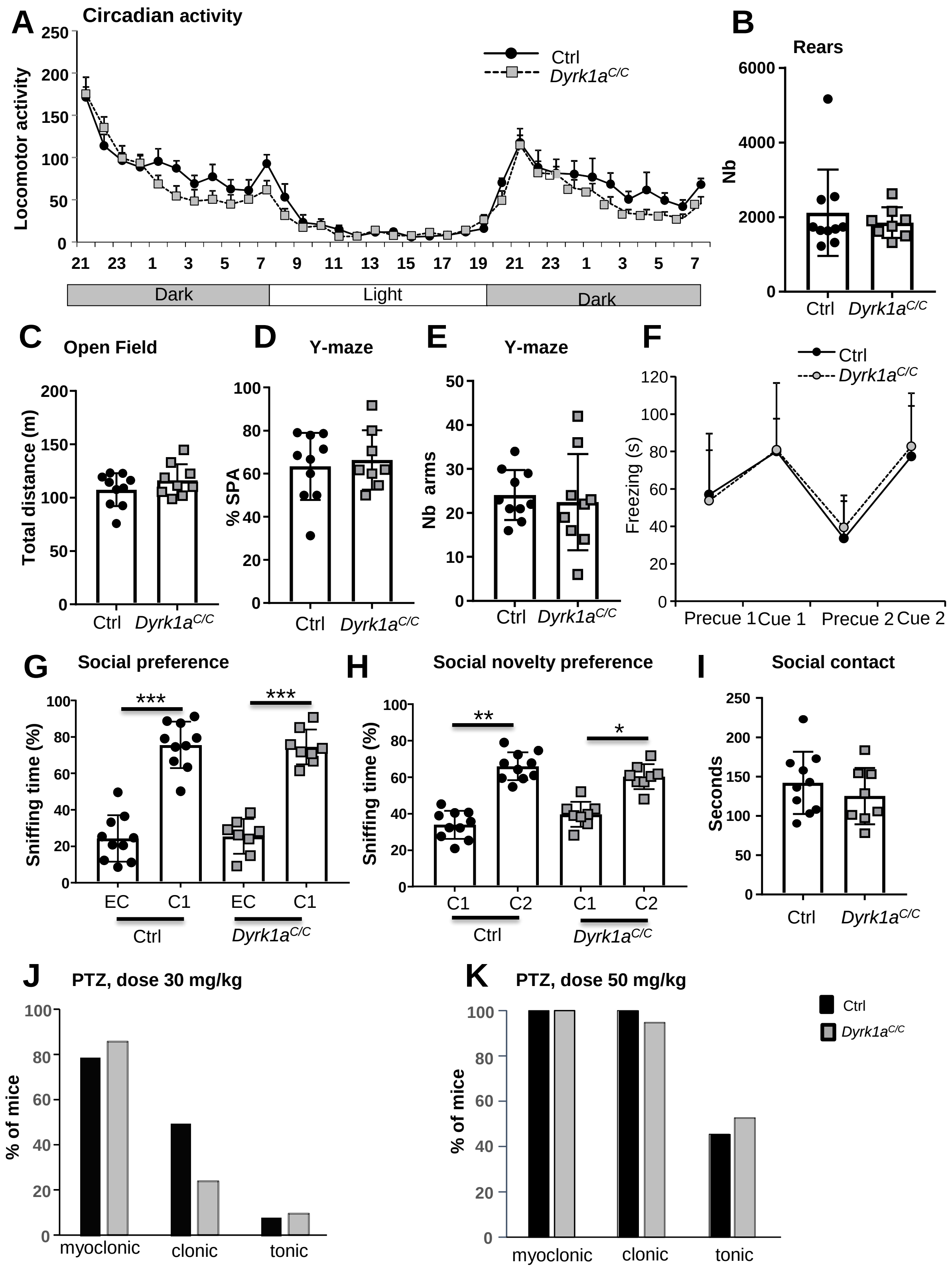


**Supplementary figure 2**

(A-B) Effects of *Dyrk1a* inactivation on circadian activity. Locomotor activity during circadian analysis was comparable between *Dyrk1a*$^{C/C}$ and control mice through the light/dark cycle.

Numbers on the Y-axis represent the hours. Data are presented as mean ± SEM for each hour. (B) Dot plot of the total number of rears registered during the whole 35H-period of circadian analysis. (C) The total distance travelled during 30 min within the OF was comparable between genotypes. (D) Working memory assessed by percentage of spontaneous alternation within the arms of the Y maze was not impacted by inactivation of Dyrk1a in *Dyrk1a*$^{C/C}$ mice. (E) The locomotor activity assessed by the number of arm entries was also similar between the two genotypes. (F) In the fear conditioning test, the baseline level of immobility (precue 1 and precue 2) and the cued freezing performances (cue 1 and cue 2) in a new context were comparable between genotypes. (G-I) Assessment of social behavior in the Crawley three-chamber test shows that both genotypes spend more time exploring the cage containing a congener than the empty cage (G; paired t-test congener vs empty cage: ctrl, ***p<0.001 and *Dyrk1a*$^{C/C}$ ***p<0.001) and exploring the novel than familiar congener (H; paired t-test new congener vs familiar congener: ctrl, **p=0.002 and *Dyrk1a*$^{C/C}$ *p=0.019). (I) Social contact assessed by measuring to time spent sniffing both congeners during the test for novelty preference was similar between mutant and control mice. (J-K) Epileptic susceptibility was tested with the injection of two doses of PTZ. Percentage of mice reaching myoclonic, clonic and tonic seizure stage were similar between the two genotypes at dose 30 mg/kg body weight (H; n=25 ctrl and n=20 *Dyrk1a*$^{C/C}$ mice) and 50 mg/kg body weight (I; n=22 ctrl and n=19 *Dyrk1a*$^{C/C}$ mice). Data are presented as point plots with mean ± SD.

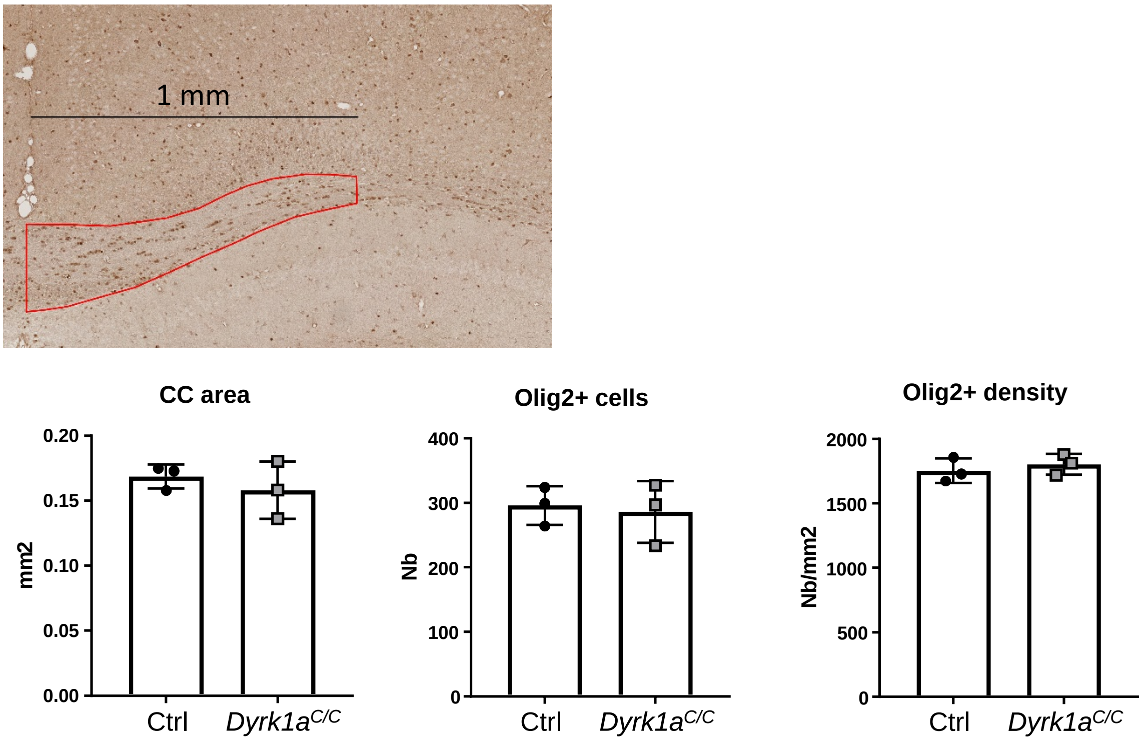


**Supplementary figure 3**

Representative image of the corpus callosum (cc) showing the selected area in which olig2+ cells were counted. A distance of 1 mm was measured and the underneath corpus callosum was selected. The cc area as well as the number of olig2+ cells and the olig2+ cell density did not differ between *Dyrk1a*$^{C/C}$ and control animals. Data are presented as point plots with mean ± SD (each dot represents the mean count of 3 serial sections).

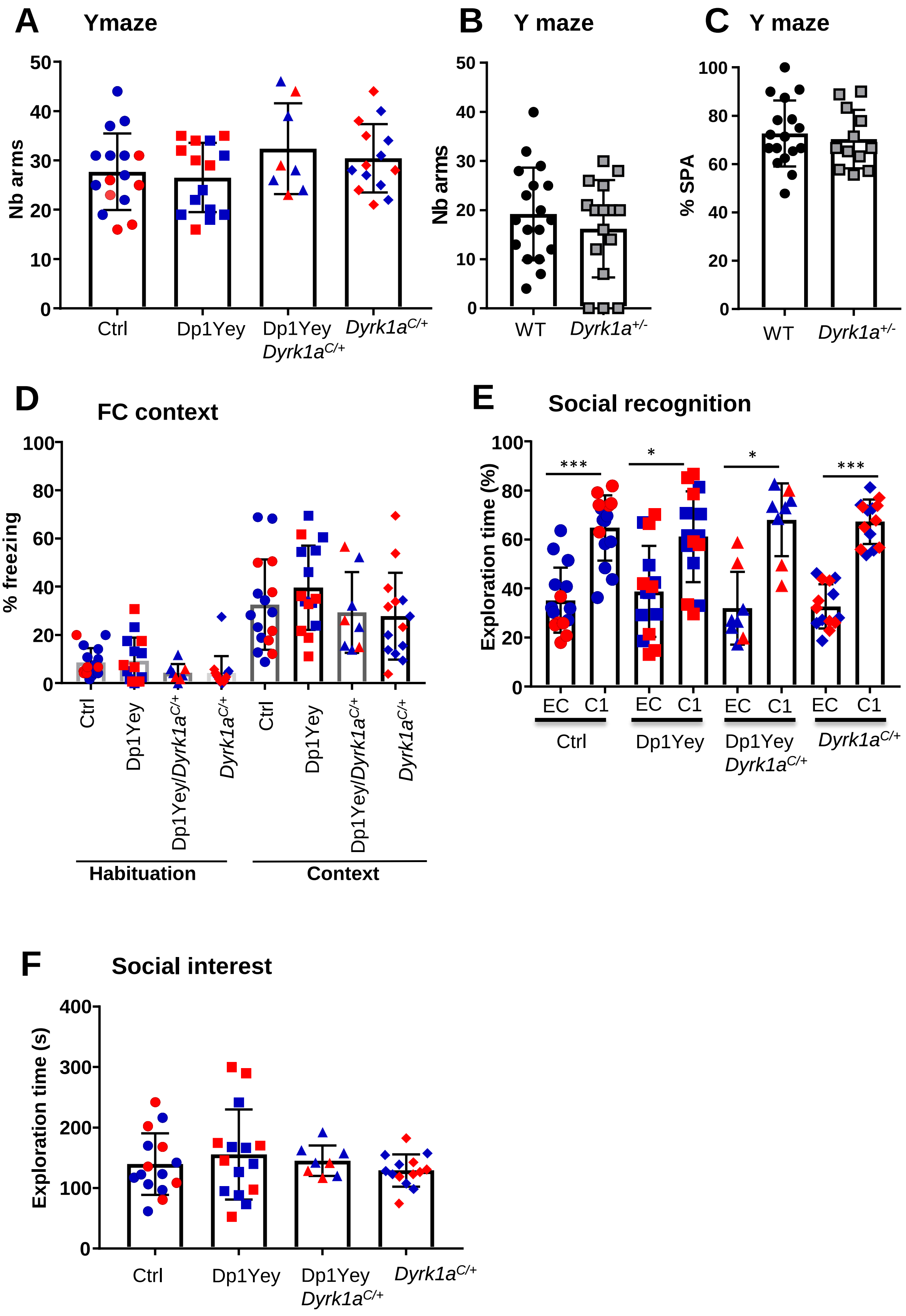
A
Ymaze
Nb arms
50
40
30
20
10
0
Ctrl
Dp1Yey
Dp1Yey
Dyrk1a C/+
Dyrk1a C/+
B
Y maze
Nb arms
WT
Dyrk1a +/-
C
Y maze
% SPA
100
80
60
40
20
0
WT
Dyrk1a +/-
D
FC context
% freezing
Ctrl
Dp1Yey
Dp1Yey/Dyrk1a C/+
Dyrk1a C/+
Habituation
Context
E
Social recognition
Exploration time (%)
***
*
*
***
EC
C1
Ctrl
Dp1Yey
Dp1Yey
Dyrk1a C/+
Dyrk1a C/+
F
Social interest
Exploration time (s)
400
300
200
100
0
Ctrl
Dp1Yey
Dp1Yey
Dyrk1a C/+
Dyrk1a C/+

**Supplementary figure 4**

(A) The locomotor activity assessed by the number of arm entries in the Y maze was similar between genotypes. (B-C) Activity (B) and working memory (C) was assessed in *Dyrk1a* full heterozygous knockout ($Dyrk1a^{+/-}$) mice in the Y maze showing no effect of *Dyrk1a* haploinsufficiency (only males were analyzed here). (D) In the fear conditioning test, the baseline level of immobility during the habituation period was similar between genotypes and contextual freezing performance in the same environment after conditioning was also comparable between genotypes. (E-F) Assessment of social behavior in the Crawley three-chamber test shows that all genotypes spend more time exploring the cage containing a congener than the empty cage (E; paired t-test congener vs empty cage: ctrl, ***p<0.001; Dp1Yey, *p=0.03; Dp1Yey/$Dyrk1a^{C/C}$ , *p=0.01; $Dyrk1a^{C/C}$, p***p<0.001). Moreover, no difference was found between genotypes in the total time spent sniffing the cage containing a congener (F). Data are represented as point plots with mean ±SD. Males are in blue and females are in red.
